# Phenotype-associated spatial biomarker discovery in spatial transcriptomics with spHOT

**DOI:** 10.64898/2026.08.11.744312

**Authors:** Hoeyoung Kim, Donghee Kim, Sangwook Jung, Sangseon Lee, Kwangsoo Kim

**Affiliations:** Interdisciplinary Program in Bioinformatics, Seoul National University, Seoul, Republic of Korea.; Biomedical Research Institute, Seoul National University Hospital Seoul, Republic of Korea.; Department of Artificial Intelligence, Inha University, Incheon, Republic of Korea.; Department of Transdisciplinary Medicine, Seoul National University Hospital, Seoul, Republic of Korea.; Department of Medicine, College of Medicine, Seoul National University, Seoul, Republic of Korea.; Center for Data Science, Healthcare AI Research Institute, Seoul National University Hospital, Seoul, Republic of Korea.

## Abstract

Spatial transcriptomics now profiles patient cohorts at single-cell resolution, enabling analysis of disease-associated cell organization in situ. However, discovering such spatial biomarkers remains challenging because relevant structures occur at unknown scales and cell-or niche-level annotations are rarely available. We present spHOT, a deep learning framework that localizes phenotype-associated spatial biomarkers from sample-level labels. spHOT combines spatial foundation model embeddings, a hierarchical domain tree for multi-resolution tissue representation, and a teacher-student multiple instance learning architecture that converts sample labels into cell-level biomarker scores. In controlled simulations and real-tissue benchmarks, spHOT outperformed existing spatial and single-cell methods in localizing ground-truth biomarkers. Across fibrotic, metabolic, and autoimmune disease datasets, spHOT recovered disease-relevant niches and tissue states reported by supervised analyses in the original studies. Cross-disease application of spHOT transferred biomarkers across chronic lung diseases without retraining. spHOT enables scalable, annotation-efficient spatial biomarker discovery in cohort-scale spatial transcriptomics.

## 1 Main

Spatial organization of cells is a fundamental determinant of tissue function and disease progression[1][2]. In complex disease tissues, pathological states are shaped not only by the molecular profiles of individual cells, but also by their spatial arrangement, local cellular neighborhoods and their coordinated interactions[3][4]. These spatial features can define key disease-associated cellular communities, or spatial biomarkers, whose pathological significance is not captured by cell type abundance or differential gene expression alone[5][6]. Spatial transcriptomics (ST), which jointly measures gene expression and cellular location within intact tissue sections, provides an opportunity to interrogate these spatial cellular communities in their native context[7][8]. Imaging-based ST platforms in particular profile gene expression at single-cell resolution, enabling characterization at the level of individual cells[9][10].

Recently, ST studies are increasingly conducted at cohort scale, profiling a large number of samples across disease conditions[11][12], and computational methods have been developed alongside this shift. Early approaches performed spatial domain detection within individual samples, integrating gene expression[13][14][15][16][17][18] or categorical cell labels[19][20] with spatial coordinates. Multi-sample frameworks then extended niche characterization across samples[21][22], and spatial foundation models have more recently learned generalizable representations of local cellular neighborhoods[23][24]. In parallel, multiple instance learning (MIL) and related weakly supervised methods have enabled cell-level disease association with sample-level labels in single-cell data[25][26][27][28]. However, most ST methods focus solely on identifying domains[13][14][15][16][17][18], or rely heavily on pre-annotated cell types for both methodology[29] and downstream interpretations of spatial biomarkers[21][22], or focus mainly on low-resolution sequencing-based platforms[30]. On the other hand, weakly supervised disease association methods typically place less emphasis on modeling spatial organization[25][26][27][28]. Consequently, current approaches remain limited in their ability to localize phenotype-associated spatial biomarkers from cohort-scale imaging-based ST data.

This limitation is compounded by several challenges. First, disease-associated spatial structures can emerge at highly variable organizational scales, ranging from small immune aggregates to large fibrotic or lymphoid niches[31], making it unclear a priori which spatial resolution is most informative. Second, the identification and evaluation of spatial biomarkers remains supervised and largely qualitative, with limited standardized metrics for quantitative assessment. Third, cell-level or niche-level annotations require expert histological review and are often unavailable or labor-intensive at cohort scale, whereas sample-level clinical labels are readily available[32]. Addressing these challenges requires searching across spatial cellular resolutions, quantifying importance localization, and learning from weak supervision.

Here, we present spHOT, an MIL-based deep learning framework for discovering phenotype-associated spatial biomarkers from cohort-scale imaging-based ST data. Taking cell-level gene expression, spatial coordinates, and sample-level clinical labels as input, spHOT infers cell-level spatial biomarker scores without requiring cell-or niche-level annotations. To capture disease-relevant organization across variable tissue scales, it couples spatial foundation model embeddings with a hierarchical domain tree that searches multi-resolution cellular communities, and employs a dual-branch teacher-student MIL architecture to score spatially coherent, phenotype-associated cell populations. We further introduce a metric-based evaluation strategy for quantifying spatial biomarker localization. Through simulation benchmarks and disease cohorts across multiple imaging-based ST platforms, we demonstrate that spHOT accurately localizes disease-associated spatial communities and recovers biologically meaningful tissue organization from sample-level labels alone.

## 2 Results

### 2.1 Identification of phenotype-associated spatial biomarkers with spHOT

spHOT localizes phenotype-associated spatial biomarkers from multi-sample ST datasets with sample-level case/control labels. Given cell-level gene expression, spatial coordinates, and binary phenotype labels, spHOT classifies sample phenotypes and produces cell-level spatial biomarker scores (cell scores) that quantify each cell’s contribution to the case/control distinction. These scores enable the spatial localization of case-associated regions within individual tissue samples.

The framework proceeds in four stages (Fig. 1). First, a pretrained spatial foundation model encodes each cell’s gene expression and local neighborhood into a low-dimensional embedding and assigns cells to initial domains[24], independent of sample labels (Fig. 1a). Second, agglomerative hierarchical clustering of domain centroids constructs a domain tree spanning coarse-to-fine tissue resolutions; these clusters are termed metadomains, representing shared tissue units across samples and phenotypes (Fig. 1b). Third, a dual-branch teacher-student MIL architecture[33] operates at each metadomain level to jointly classify samples and infer case-associated cells (Fig. 1c). The teacher branch computes metadomain-level case association scores, while the student branch resolves case-associated cells at single-cell resolution using metadomain-aware sampling. The two branches share an encoder and are trained via alternating optimization. Finally, spHOT selects the metadomain level that maximizes spatial coherence of cell scores, using a composite spatial autocorrelation score based on Moran’s I[34] and Geary’s C[35] (Fig. 1d). (*k*) maximizing the composite spatial score is selected for spatial biomarker discovery. **e** Downstream analytical framework of spHOT for the characterization and discovery of disease-associated spatial biomarkers.

**Fig. 1:**
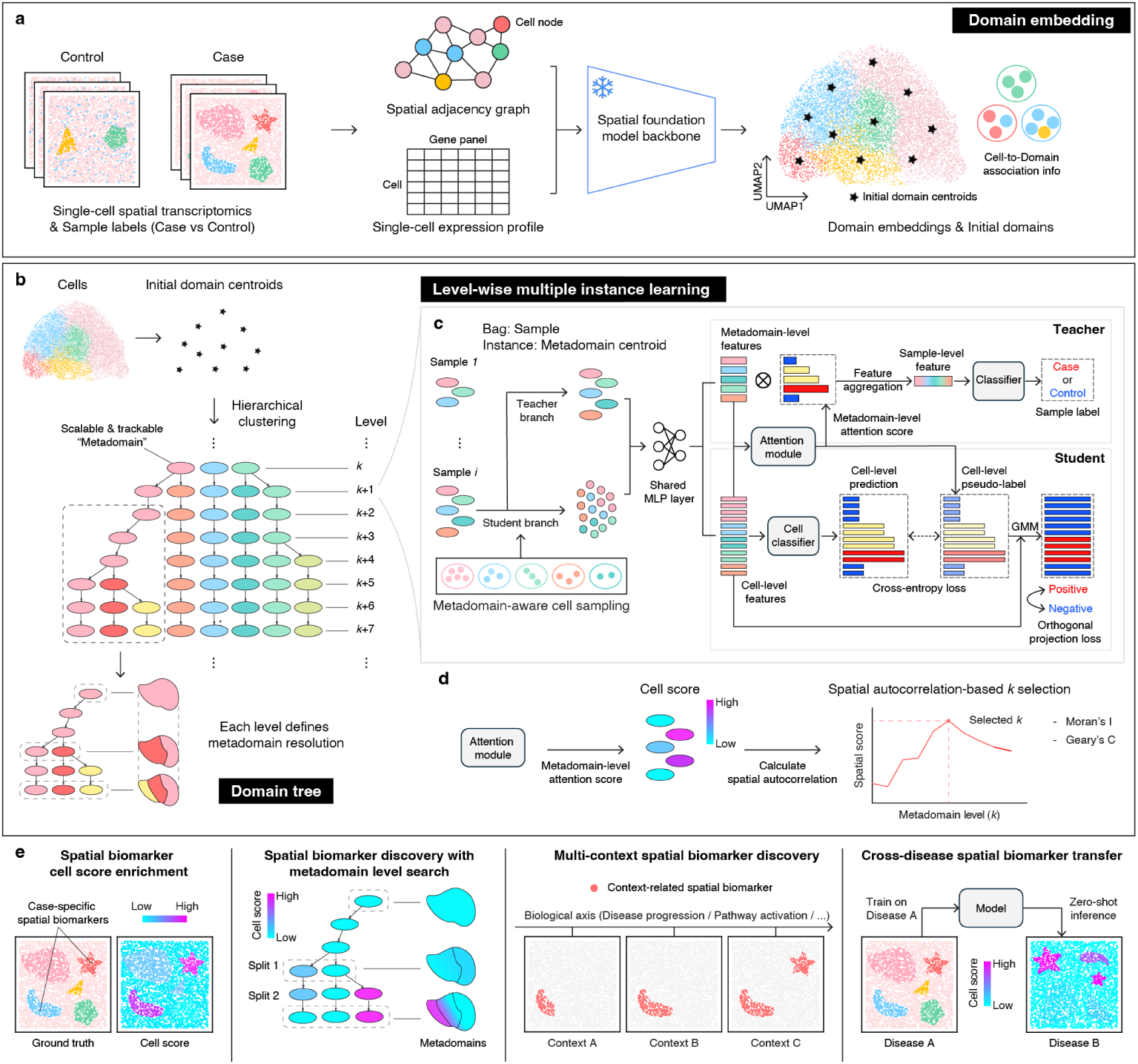
Overview of spHOT. a–d. spHOT is a modular framework proceeding in four stages: Domain embedding (**a**); Domain tree (**b**); Level-wise multiple instance learning (**c**); Spatial autocorrelation-guided level selection (**d**). **a** Imaging-based, single-cell spatial transcriptomics data from case and control samples are processed through a spatial foundation model backbone in a zero-shot manner. A spatial adjacency graph built from cell coordinates is combined with expression profiles as input. The model produces low-dimensional domain embeddings and assigns each cell to an initial domain via learned prototype vectors. **b** Initial domain centroids are hierarchically clustered into a domain tree, in which each level (*k*) defines a distinct metadomain resolution. **c** At each metadomain resolution, a dual-branch teacher-student MIL architecture is applied, with both branches sharing a common MLP encoder. The teacher branch treats each sample as a bag of metadomain centroids, computes metadomain-level attention scores, and predicts sample labels (case or control). The student branch operates on cells drawn by metadomain-aware sampling, produces cell-level predictions, and generates cell-level pseudo-labels using a Gaussian mixture model (GMM). The student is trained with cross-entropy loss on pseudo-labels and orthogonal projection loss (OPL) to separate positive (case-specific) and negative (control-like) cell representations. **d** Metadomain-level attention scores from the teacher branch are propagated to cells, and spatial autocorrelation is computed on cell scores using Moran’s I and Geary’s C at each level (*k*), combined into a single composite spatial score. The level

Beyond cell score inference, spHOT provides an integrated downstream analysis framework for evaluating and interpreting phenotype-associated spatial organizations (Fig. 1e). This includes enrichment scoring for quantitative assessment of spatial biomarker localization, multi-scale domain tree analysis for characterizing scale-dependent spatial biomarkers, multi-context comparison for identifying disease-subtype-specific spatial organization, and cross-disease transfer analysis for evaluating shared spatial biomarkers across disease cohorts.

### 2.2 spHOT outperforms existing methods in spatial biomarker discovery under controlled simulation

To evaluate spHOT and existing methods for spatial biomarker localization, we constructed controlled simulation datasets in which spatial arrangement, cell type composition, and gene expression perturbation could be varied independently. Realistic single-cell expression profiles were generated using scCube[36] trained on a real-world CosMx dataset[37], and seven anonymized cell types were placed in 100 *×* 100 pixel fields to create synthetic ST samples (Fig. 2a,b and Methods). Control samples contained randomly mixed cell types with equal compositions and were annotated as background. In case samples, ground truth spatial biomarkers were generated by modifying the spatial arrangement, composition, or transcriptional state of selected cell populations according to each scenario, as described below. Each scenario included 20 control and 20 case samples and was evaluated using five-fold cross-validation, with cell scores computed for each held-out test sample exactly once (Methods).

**Fig. 2:**
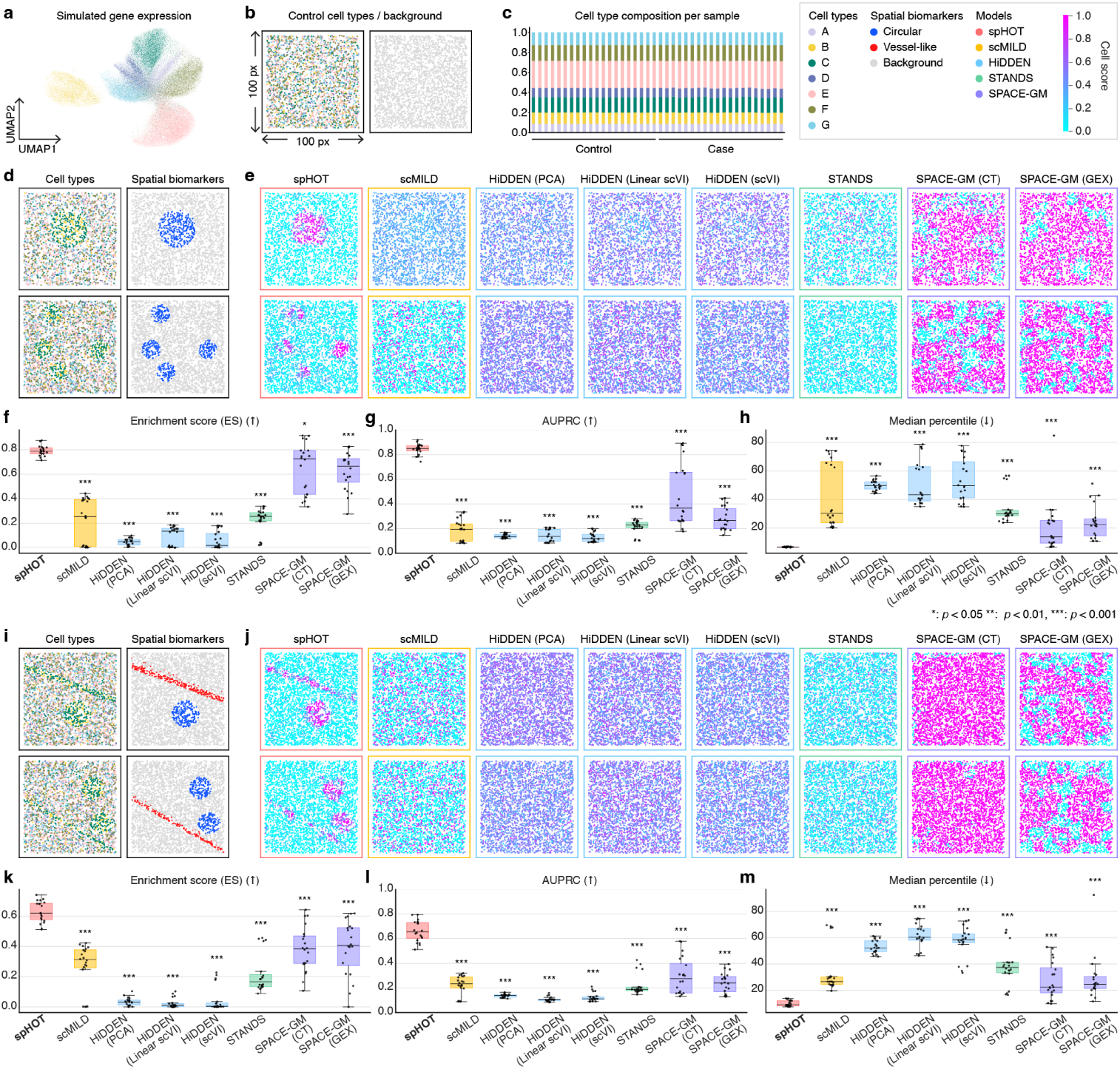
Performance evaluation of spatial biomarker discovery on simulated datasets. **a** UMAP visualization of simulated gene expression profiles generated using scCube, comprising seven anonymized cell types (A–G). **b** Representative spatial layouts of a control sample, colored by cell type (left) and ground truth spatial biomarkers (right), showing a random distribution of all cell types. **c** Cell type composition per sample across all control and case samples. **d**–**h** Evaluation on the circular-only spatial biomarker scenario. **d** Two representative spatial layouts, colored by cell type (left) and ground truth spatial biomarker (right). **e** Continuous heatmaps of normalized cell scores within [0, 1] range for all methods. Box outlines indicate method type. Upper and lower rows correspond to the same representative samples as in **d**. **f** –**h** Bench-marking performance across 20 test-set case samples, showing **f** enrichment score (ES); **g** area under the precision-recall curve (AUPRC); **h** median ground truth percentile. **i**–**m** Evaluation on the mixed circular and vessel-like spatial biomarker scenario. Panels **i**, **j** and **k**–**m** are formatted identically to **d**, **e**, and **f** –**h**, respectively. Box plots show the median (centerline), interquartile range (box), and 1.5*×* IQR (whiskers); each point is one sample. The best-performing method is highlighted in bold. Asterisks denote significant differences from the best-performing method (paired Wilcoxon signed-rank test; \**p <* 0.05, \*\**p <* 0.01, \*\*\**p <* 0.001).

We benchmarked spHOT against seven types of recent methodologies that produce cell-level importance scores, including spatial approaches that incorporate cell coordinates (SPACE-GM[29] and STANDS[30]) and single-cell approaches that rely on expression alone (scMILD[28] and HiDDEN[25]). Performance was evaluated based on the ability to localize ground truth spatial biomarker cells using enrichment score (ES), area under the precision-recall curve (AUPRC), and median ground truth percentile, with additional metrics (Methods).

The first three scenarios tested spatial-only biomarkers, in which case and control samples had matched cell type composition and gene expression distributions but differed in the spatial organization of selected cell types (Fig. 2c). These scenarios included compact circular aggregates (Fig. 2d), elongated vessel-like structures (Extended Data Fig. 1a), and samples containing both morphologies (Fig. 2i). The next two introduced non-spatial signals on top of a circular spatial biomarker, creating confounding scenarios. Circular and composition altered (CC) biomarkers included a case-associated increase in the abundance of biomarker-forming cell types (Extended Data Fig. 2c,d), whereas circular and composition altered with perturbation (CCP) biomarkers additionally incorporated a transcriptionally perturbed stressed cell state (F*) (Extended Data Fig. 2k,l).

In the spatial-only settings, spHOT consistently localized ground truth biomarker structures across different spatial morphologies. For circular biomarkers, spHOT achieved the highest ES, AUPRC, and lowest median percentile, outperforming all other methods (Fig. 2e–h, Extended Data Fig. 3a,b, and Supplementary Table 2–3). When circular and vessel-like structures coexisted within the same sample, spHOT maintained its advantage (Fig. 2j–m, Extended Data Fig. 3c,d, and Supplementary Table 4–5), simultaneously recovering multiple morphologies. In the vessel-like-only setting, spHOT again showed strong performance with fewer false positives (Extended Data Fig. 1b–e, Extended Data Fig. 3e,f, and Supplementary Table 6–7). In the spatial-only settings, three spatial methods consistently outperformed single-cell methods, confirming that spatial organization constitutes an axis of biological information orthogonal to gene expression.

In the CC scenario, spHOT outperformed other methods by preferentially localizing biomarkers (Extended Data Fig. 2e), while single-cell methods improved but often assigned high scores broadly across the expanded cell type, including cells outside the biomarker regions (Extended Data Fig. 2e–h, Extended Data Fig. 3g,h, and Supplementary Table 8–9). In the CCP scenario, expression-based methods improved further, whereas spHOT remained among the top-performing methods, maintaining accurate localization (Extended Data Fig. 2m–p, Extended Data Fig. 3i,j, and Supplementary Table 10–11).

Beyond biomarker localization, we assessed phenotype classification performance from the same learned representations. We included two existing supervised single-cell methods: ProtoCell4P[38] and Hier-MIL[26]. Sample-level classification AUROC tracked the cell score trends, and spHOT achieved the highest or second-highest AUROC across all scenarios (Extended Data Table. 1).

### 2.3 spHOT accurately detects clinician-annotated immune hotspots in idiopathic pulmonary fibrosis tissue

Having established spHOT’s performance on simulated datasets, we next applied it to real disease tissues, utilizing the Xenium idiopathic pulmonary fibrosis (IPF) dataset[39]. Ground truth spatial biomarkers were focused on two immune hotspot niches: tertiary lymphoid structures (TLS) and mixed inflammation aggregates. spHOT achieved robust sample classification across five folds (mean AUROC = 0.933, s.d. = 0.149), indicating cell scores inferred from discriminative case/control signals.

For TLS localization, spHOT most consistently concentrated high cell scores within annotated TLS regions, achieving the highest enrichment (Fig. 3a–e, Extended Data Fig. 4a,b). Because only two samples contained TLS annotations, we did not perform pairwise statistical testing. By contrast, competing methods either produced diffuse high-score patterns or prioritized areas outside of the annotation (Fig. 3b).

**Fig. 3:**
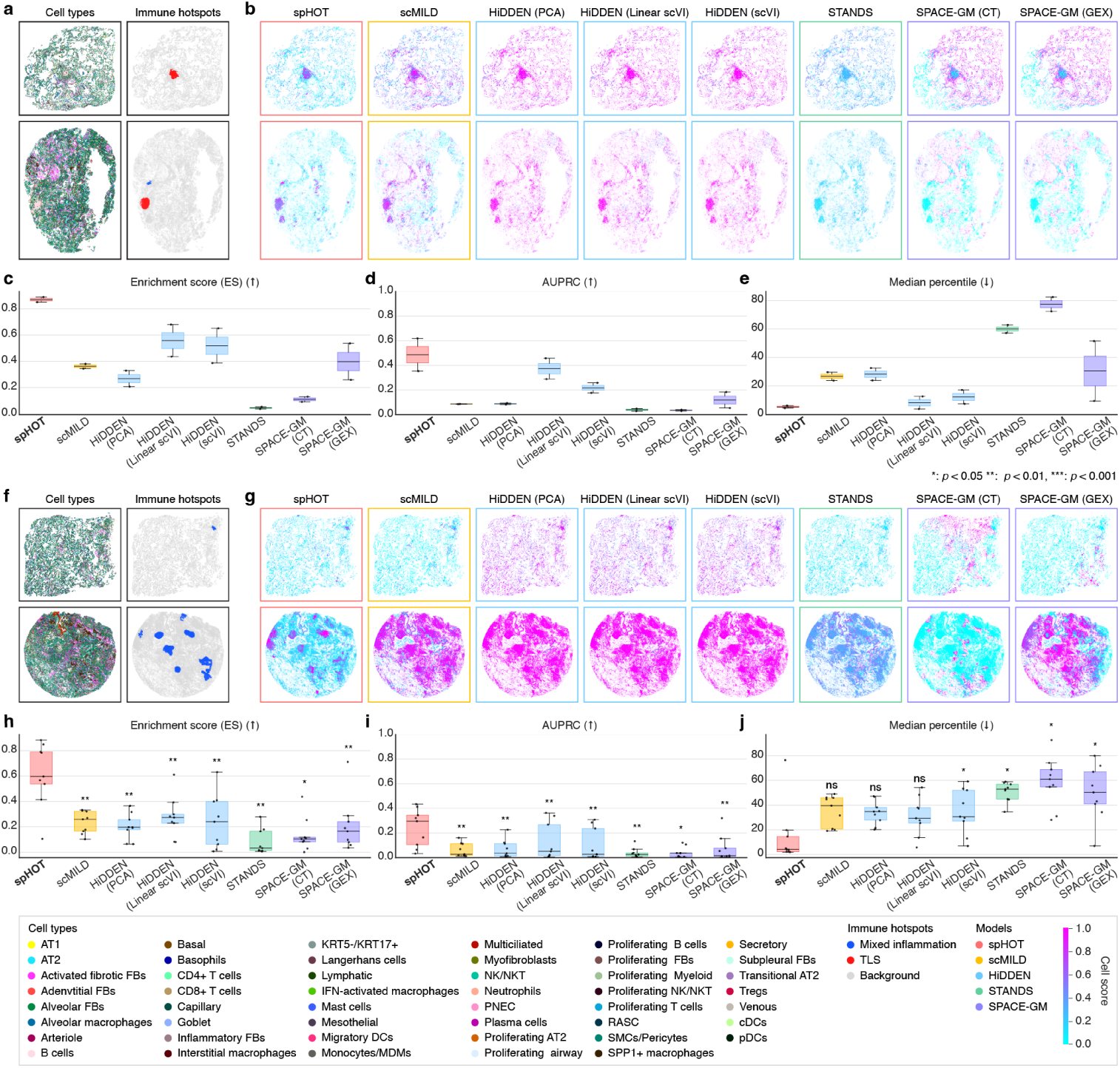
Benchmarking spHOT and existing methods for immune hotspot discovery in Xenium idiopathic pulmonary fibrosis. **a**–**e** Evaluation on tertiary lymphoid structures (TLS). **a** Two representative samples showing cell type annotations (left) and ground truth TLS regions (right). **b** Continuous heatmaps of normalized cell scores within [0, 1] range for all methods. Box outlines indicate method type. Upper and lower rows correspond to the same representative samples as in **a**. **c**–**e** Benchmarking performance on TLS ground truth discovery across 2 test-set case samples. Box plots show **c** ES; **d** AUPRC; **e** median ground truth percentile. **f** –**j** Evaluation on mixed inflammation. Panels **f**, **g** and **h**–**j** are formatted identically to **a**, **b**, and **c**–**e**, respectively. **h**–**j** Benchmarking performance on mixed inflammation ground truth detection across 9 test-set case samples. Box plots show the median (center-line), interquartile range (box), and 1.5*×* IQR (whiskers); each point is one sample. The best-performing method is highlighted in bold. Asterisks denote significant differences from the best-performing method (paired Wilcoxon signed-rank test; \**p <* 0.05, \*\**p <* 0.01, \*\*\**p <* 0.001, ns, not significant).

For mixed inflammation localization, spHOT separated more clearly from competing methods and achieved significantly higher performance (Fig. 3f–j, Extended Data Fig. 4c,d, and Supplementary Table 12–13). Mixed inflammation regions represent heterogeneous immune aggregates that can vary substantially in size, density, and spatial dispersion, making them difficult to identify from cell type composition alone[31]. Across representative samples spanning both rare, focal hotspots and abundant, broadly distributed inflammation (Fig. 3f), spHOT preferentially assigned high scores to the clinician-annotated regions, suggesting that metadomain-wise cell scores capture spatially organized immune pathology rather than immune cell abundance or diffuse inflammatory signal. Together, these results show that spHOT localizes spatially organized immune structures in fibrotic lung tissue directly from sample-level labels.

### 2.4 Hierarchical metadomain framework of spHOT enables de novo discovery of a profibrotic macrophage niche in IPF

Beyond detecting pre-annotated immune hotspots, we next asked whether spHOT could identify unannotated, biologically relevant spatial biomarkers de novo. In the selected metadomain resolution (*k* = 21) of the Xenium IPF dataset, the high-scoring regions were not limited to the immune hotspots but were also localized to unannotated airway-associated niches. To further explore them, we traced the corresponding metadomain lineages through the hierarchical domain tree (Fig. 4a).

**Fig. 4:**
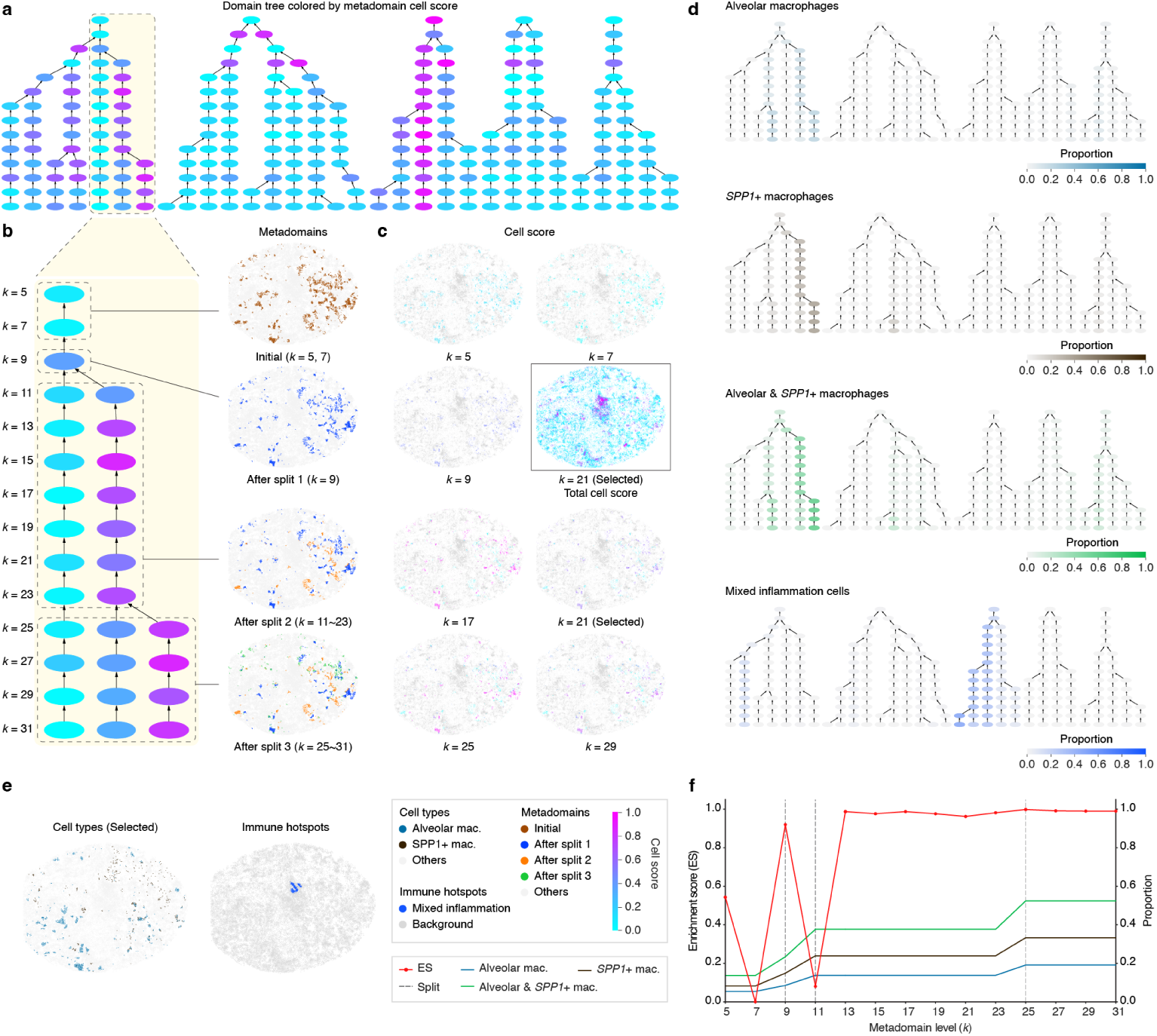
De novo spatial biomarker discovery via hierarchical metadomain level search in IPF tissue. **a** Domain tree visualization for test-set IPF samples in a single representative cross-validation fold with tree nodes colored by metadomain-level cell score. Tree spans from *k* = 5 (top) to *k* = 31 (bottom), with two metadomains increment per level. The tinted dashed box highlights the lineage of the discovered macrophage niches, zoomed-in in **b**. **b** Spatial visualization of the high-scoring lineage across successive tree splits in representative sample VUILD91MA. Column of tissue map shows metadomain assignments at four representative levels, with the tracked lineage metadomain highlighted and all others shown in grey. **c** Cell score spatial visualizations at selected metadomain resolutions (*k* = 5, 7, 9, 17, 21, 25, 29). Scores are normalized within [0, 1] range and displayed as a continuous heatmap. The boxed panel highlights *k* = 21 as the spatial autocorrelation-selected resolution, showing the total cell score map across the full tissue section. **d** Cell type proportion projections onto the metadomain tree for four different categories: alveolar macrophages, *SPP1* + macrophages, combined alveolar and *SPP1* + macrophages, and mixed inflammation-annotated cells. Node color intensity reflects the proportion of each category within each metadomain at each level, overlaid on the same tree structure as in **a**. **e** Spatial visualization of selected macrophage cell types (left) and immune hotspot annotations (right). **f** Averaged ES (red, left axis) and macrophage cell type proportions (right axis) across metadomain resolutions *k* = 5 to *k* = 31, averaged across test-set IPF samples. Alveolar macrophage, *SPP1* + macrophage, and combined alveolar and *SPP1* + macrophage proportions within the high-scoring metadomain lineage are shown. Dashed vertical lines indicate tree split events.

At coarse resolutions (*k* = 5, 7), niche regions were merged into broad, heterogeneous metadomains, diluting their disease-associated signal. After the first splits at *k* = 9, 11, the niches separated from the surrounding tissue and the high cell scores concentrated in a more aggregated form enriched for macrophages. This localization persisted at *k* = 21 (Fig. 4a–c and Extended Data Fig. 5a–n). As shown in Fig. 4d, cell type proportion overlays on the domain tree showed that the high-scoring lineage was enriched for alveolar macrophages and *SPP1* + macrophages, consistent with the airway-accumulated macrophage niche identified by Vannan et al.[39] through supervised niche analysis. In contrast, immune hotspots mapped to distinct tissue locations and separate metadomain lineages, indicating that the macrophage niche represents an independent spatial biomarker lineage rather than a redundant observation (Fig. 4d,e and Extended Data Fig. 5o).

To quantify the resolution dependence of this discovery, we computed macrophage niche detection ES across metadomain levels, averaged over IPF samples (Fig. 4f and Methods). While ES was unstable at coarse resolutions, it increased sharply after *k* = 9, 11 and stabilized through *k* = 21 and beyond. The increasing proportion of both alveolar and *SPP1* + macrophages within the metadomain lineage mirrored this stabilization pattern of the ES. These results show that the domain tree is not merely a computational convenience: overly coarse resolutions obscure the niche by merging it with unrelated domains, whereas appropriate resolutions reveal a coherent disease-associated spatial structure. This multi-resolution traversal enables spHOT to discover spatial biomarkers whose organizational scale is unknown a priori.

This de novo discovery converges on a central finding of Vannan et al., who described progressive accumulation of *FABP4* + alveolar and *SPP1* + macrophages within airspaces as a late event in alveolar remodeling, following capillary and AT1 loss, epithelial transition, and fibroblast activation[39]. Because *SPP1* + macrophages are reported to be enriched within and around fibroblastic foci in IPF and are associated with myofibroblast activation and osteopontin deposition in the extracellular matrix (ECM)[40][41], this macrophage niche represents a plausible profibrotic spatial biomarker rather than a passive immune accumulation. spHOT recovered this niche without using macrophage-specific prior knowledge or niche-level annotations, demonstrating its ability to identify biologically meaningful spatial organization de novo.

### 2.5 Each architectural component of spHOT contributes to the discovery of spatial biomarkers

spHOT integrates three key components to translate sample labels into spatially coherent cell scores: a hierarchical domain tree, metadomain-aware cell sampling, and a teacher-student MIL architecture. To assess their individual contributions, we performed three ablation studies on simulated datasets and the Xenium IPF dataset, removing the domain tree (Ablation 1), cell sampling (Ablation 2), or the student branch (Ablation 3) in turn. Removing the domain tree localization reduced stability, particularly in simulations, confirming the benefit of hierarchical resolution search (Extended Data Fig. 6c–h, and Supplementary Table 14–17). Removing cell sampling degraded performance while substantially increasing training time (Extended Data Fig. 6a–b), indicating a role beyond efficiency in balancing cell representation (Extended Data Fig. 6c–h, i–n, and Supplementary Table 14–17). Removing the student branch particularly reduced performance in real data, reflecting the importance of cell-level refinement in complex tissues (Extended Data Fig. 6i–n, and Supplementary Table 18–19). Together, these results demonstrate that all three components contribute to accurate and efficient spatial biomarker localization.

### 2.6 spHOT identifies active and chronic diffuse tubular injury states in CosMx-profiled diabetic kidney disease

To further test whether spHOT generalizes across ST platforms and diverse tissues, we applied it to a CosMx diabetic kidney disease (DKD) dataset[42]. Spatial biomarker regions were defined as tubular injury areas spanning *HAVCR1* + injured proximal tubules (Extended Data Fig. 7a and Methods). Despite moderate sample-level classification (mean AUROC = 0.700, s.d. = 0.216), spHOT retained reliable localization of annotated injury regions, achieving the best or comparable performance across metrics (Extended Data Fig. 7b–e, Extended Data Fig. 4e,f, and Supplementary Table 20–21). Although DKD tubular injury appeared as spatially dispersed proximal tubule damage interspersed with intact nephron structures unlike the focal immune hotspots in IPF, spHOT maintained its localization ability and showed applicability across both platform type and spatial biomarker morphology.

Beyond localization, marker gene analysis revealed that high-scoring metadomains captured distinct tubular injury states rather than a single homogeneous region (Extended Data Fig. 7f and Methods). Metadomains 3, 2, 1, 5, and 7 showed elevated *IL32* with relatively low *COL4A1* and *COL4A2* expression, consistent with active tubular injury. In contrast, metadomains 6 and 4 co-expressed *IL32* together with higher collagen expression, suggesting a transition toward chronic fibrotic injury[42]. Cell type composition of metadomains further supported this interpretation: high-scoring metadomains were enriched for *HAVCR1* + injured proximal tubule cells and macrophages, whereas low-scoring metadomains showed elevated composition of *HAVCR1 −* healthy proximal tubules (Extended Data Fig. 7g). Thus, metadomain-level characterization further resolves active and chronic injury states without additional region-level annotations.

### 2.7 Multi-context comparison identifies late stage ANCA-GN spatial biomarkers resolving into functionally distinct tissue zones

We next aimed to demonstrate spHOT’s capacity for context-dependent spatial biomarker discovery. We utilized a Xenium rapidly progressive glomerulonephritis (RPGN) dataset[43], composed of multiple disease subtypes with known progression axis. Rather than performing a single case/control comparison, we used two complementary comparison settings on the same anti-neutrophil cytoplasmic antibody-associated glomerulonephritis (ANCA-GN) subtype (Fig. 5a). Setting 1 (S1; healthy control vs. ANCA-GN) was designed to identify general ANCA-GN-associated regions compared to healthy control, whereas Setting 2 (S2; systemic lupus erythematosus-associated glomerulonephritis (SLE-GN) vs. ANCA-GN) was designed to define regions in ANCA-GN relative to a milder RPGN subtype. spHOT robustly classified sample phenotypes for each setting (mean AUROC = 1.000, s.d. = 0.000 for S1 and mean AUROC = 0.785, s.d. = 0.147 for S2). Cell scores from each setting were binarized into high-and low-score groups (Methods).

**Fig. 5:**
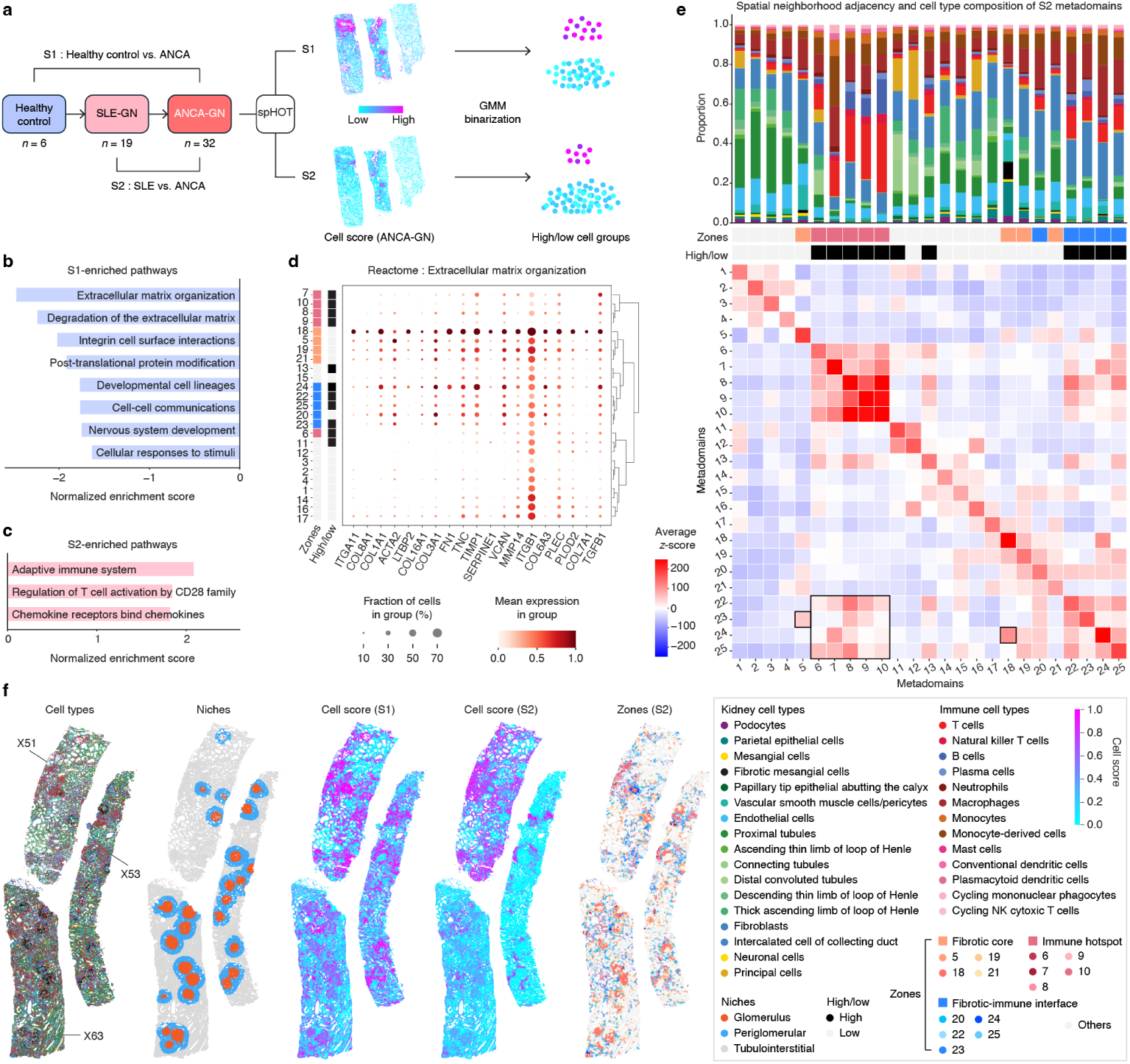
Multi-context spatial biomarker discovery identifies ANCA-GN-specific tissue zones in rapidly progressive glomerulonephritis. **a** Workflow of the multi-context comparison framework. spHOT is applied to 32 ANCA-GN samples under two independent comparison settings: S1 (Healthy control vs. ANCA-GN) and S2 (SLE-GN vs. ANCA-GN). Cell scores from each setting are binarized into high and low groups via a two-component Gaussian mixture model. **b**–**c** Reactome pathway enrichment results for S1-high-enriched pathways (**b**) and S2-high-enriched pathways (**c**), derived from preranked GSEA on pseudobulk differential expression between S1-high and S2-high cells. Bar plot shows normalized enrichment scores (NES) for top 10 significantly enriched pathways (FDR *<* 0.05). **d** Dot plot of Reactome ECM organization pathway genes across 25 S2 metadomains, hierarchically clustered by expression profile. Dot size indicates the fraction of cells expressing each gene; color intensity indicates normalized mean expression. Left axis annotations indicate zone assignment and high/low metadomain classification. **e** Top: cell type composition of all 25 S2 metadomains displayed as stacked bar plots, with zone and high/low annotations indicated below. Bottom: spatial neighborhood adjacency heatmap of S2 metadomains, displayed as average z-scores of co-occurrence frequency across all 32 ANCA-GN samples. Boxed regions highlight the fibrotic-immune interface metadomains. Color bars indicate zone assignment. **f** Spatial visualizations of three representative ANCA-GN samples. From left to right: cell type annotations; niche annotations; S1 cell scores; S2 cell scores; and S2 zone assignments. Scores are normalized to [0, 1] range and displayed as a continuous heatmap.

To characterize the molecular programs captured by each comparison, we performed pseudobulk differential expression analysis between S1-high and S2-high cells within glomerular and periglomerular regions, followed by preranked pathway enrichment analysis (Fig. 5b,c, Extended Data Fig. 8a–c, Supplementary Table 22–24 and Methods). S1-high cells were enriched for ECM organization and tissue-remodeling programs (Fig. 5b and Extended Data Fig. 8a), whereas S2-high cells were enriched for adaptive immune programs (Fig. 5c and Extended Data Fig. 8b).

We then examined the Reactome pathway genes across the 25 S2 metadomains and found S2-high signal resolved into distinct metadomain groups (Fig. 5d and Extended Data Fig. 8d). Further combining marker expression, cell type composition, and spatial neighborhood adjacency analysis revealed three major zones (Fig. 5d,e and Methods): The fibrotic core was enriched for fibrotic mesangial cells, fibroblasts, and ECM-related expression, consistent with established glomerulosclerotic regions. The immune hotspot was enriched for lymphoid and myeloid immune populations, consistent with periglomerular immune infiltration [43]. Between these regions, spHOT identified metadomains that were spatially adjacent to both the fibrotic core and immune hotspots, expressed both ECM and adaptive immune programs, but lacked the strong fibrotic mesangial cell enrichment (Fig. 5e). Therefore, we termed this zone the fibrotic-immune interface.

The high-scoring cells showed a distinct pattern between settings. As shown in Fig. 5f, representative ANCA-GN samples from different cross-validation folds showed consistent zonation and scoring patterns. In S1, all three zones were highlighted, reflecting broad disease-association relative to healthy control. In S2, however, high cell scores were selectively retained in the immune hotspot and fibrotic-immune interface, whereas the fibrotic core was attenuated. This attenuation aligned to the pathway enrichment result, suggesting that established fibrosis is shared between SLE-GN and ANCA-GN, while the immune hotspot better captures late-stage ANCA-GN-specific pathology. The fibrotic-immune interface corresponds to the spatial manifestation of the temporal transition identified by Sultana et al., from early PDGF-driven parietal epithelial cell (PEC) proliferation in inflammatory crescents to late adaptive immunity-driven, TGFβ-mediated glomerulosclerosis in fibrous crescents [43]. Thus, these results demonstrate that spHOT can be used in a multi-context manner to pinpoint functionally distinct tissue zones associated with disease progression.

### 2.8 Cross-disease transfer of spHOT reveals conserved immune hotspots that scale with COPD severity

Having established spHOT’s performance within individual disease cohorts, we continued on to ask whether spatial biomarkers learned from one disease could be cross-applied to another. The immune hotspots and profibrotic macrophage niches identified by spHOT in IPF suggested that organized immune microenvironments may represent a conserved architectural response to chronic lung injury. We therefore applied the IPF-trained spHOT model to an external Xenium chronic obstructive pulmonary disease (COPD) cohort (Fig. 6a)[44]. Despite few shared gene panels between the IPF and COPD datasets, zero-shot foundation model embeddings from the two cohorts occupied a largely overlapping latent space (Fig. 6b). The spHOT teacher branch trained on IPF was applied directly to COPD cells without retraining the model. Inferred cell scores were binarized into high-and low-score groups (Methods). High-scoring cells formed spatially discrete foci rather than diffuse signals across tissue (Fig. 6c). Pseudobulk differential expression followed by pathway enrichment analysis showed that high-scoring cells were enriched for immune-related programs (Extended Data Fig. 9a–c and Supplementary Table 25–27) compared to low-scoring cells. Cell type composition analysis further showed significant enrichment of B cells, T cells, natural killer T cells, and alveolar macrophages among high-scoring regions, while low-scoring regions were dominated by parenchymal populations and fibroblasts (Fig. 6d and Extended Data Fig. 9d). Thus, the transferred IPF model recapitulated TLS-like lymphoid aggregates and macrophage-associated immune structures in COPD without COPD-specific supervision.

**Fig. 6:**
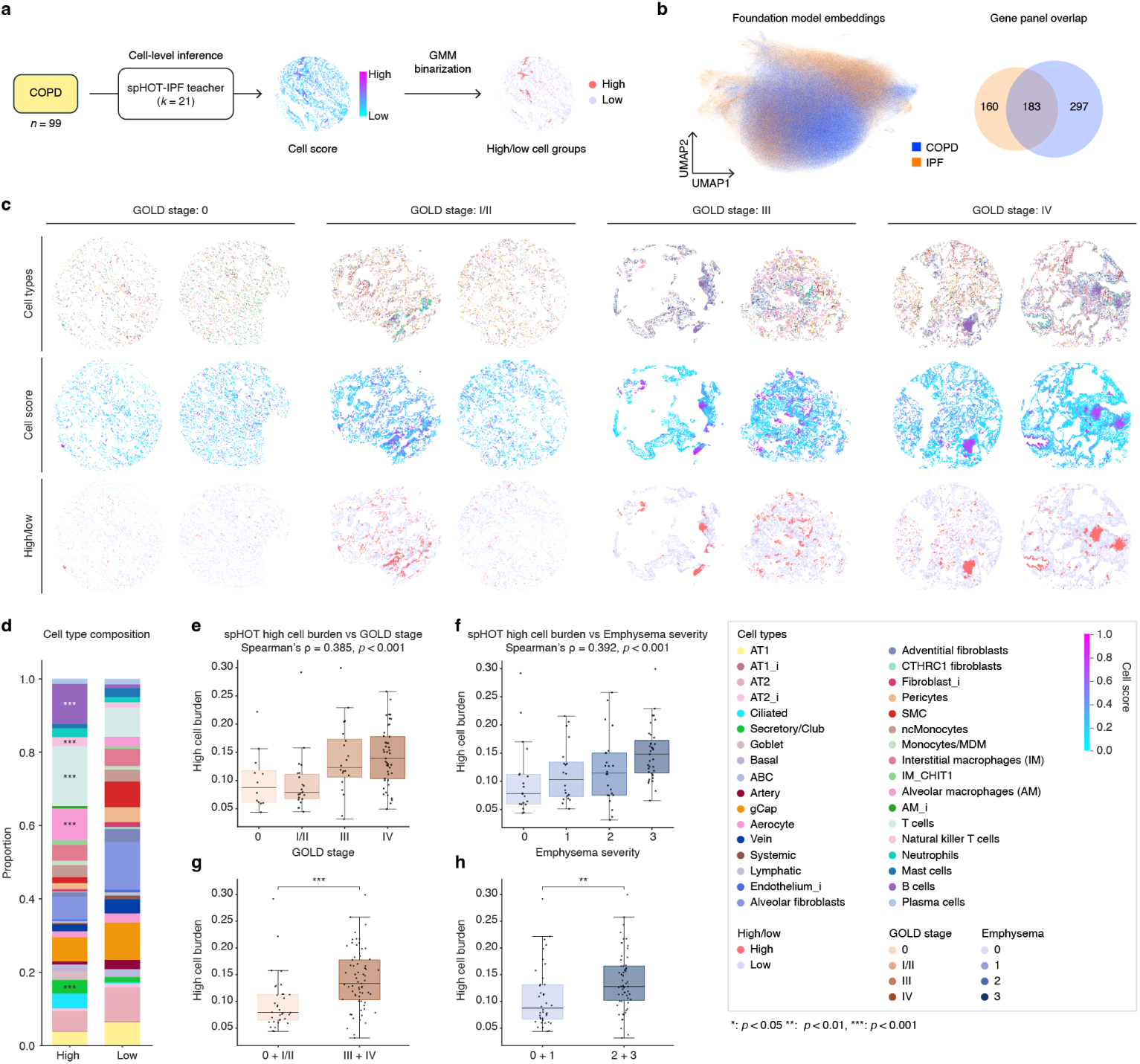
Cross-disease application of spHOT to an external cohort. **a** Workflow of the cross-disease application. The spHOT teacher trained on Xenium IPF data at *k* = 21 was applied directly to 99 Xenium COPD cores to perform cell-level attention score inference. Per-cell scores were binarized into high and low groups using a two-component Gaussian mixture model. **b** Left, UMAP of foundation model embeddings for IPF (orange) and COPD (blue) cells. Right, Venn diagram of gene panels profiled in each dataset. **c** Xenium COPD cores across GOLD stages 0, I/II, III, and IV. Two representative cores are shown per stage. Rows show cells colored by cell type annotations (top), continuous spHOT cell scores (middle), and binarized high/low groups (bottom). Scores are normalized within [0, 1] range and displayed as a continuous heatmap. **d** Cell type composition of high versus low cells, aggregated across all cores. Asterisks denote cell types significantly enriched in the high group (*p*-value calculated with Mann-Whitney U test on per-core proportions, Benjamini-Hochberg corrected). **e**,**f** Per-core high cell burden by GOLD stage (**e**) and radiographic emphysema severity (**f**). Spearman correlation and significance is shown. **g**,**h** High cell burden grouped by GOLD stage into mild (0 + I/II) versus severe (III + IV) disease (**g**) and by emphysema severity into low (0 + 1) versus high (2 + 3) (**h**). Two-sided Mann-Whitney U test is used to assess significance. Box plots show the median (centerline), interquartile range (box), and 1.5*×* IQR (whiskers); each point is one core (*n* = 99). Significance: \**p <* 0.05, \*\**p <* 0.01, \*\*\**p <* 0.001.

We then asked whether the burden of transferred high-scoring cells reflected disease severity. High-cell burden, defined as the fraction of high-scoring cells per core, increased with both GOLD stage (Spearman’s *ρ* = 0.385, *p <* 0.001; Fig. 6e) and radio-graphic emphysema severity (*ρ* = 0.392, *p <* 0.001; Fig. 6f). Dichotomized comparisons confirmed that severe COPD cores carried greater high-cell burden than control or mild disease cores, both by functional staging (GOLD III/IV versus 0/I/II, *p <* 0.001; Fig. 6g) and by emphysema severity (2/3 versus 0/1, *p <* 0.01; Fig. 6h) (Supplementary Table 28). These findings are consistent with the original study, which reported expansion of inflammatory, remodeling, and immune niches with increasing disease severity[44]. Together, these results show that spHOT trained on fibrotic lung disease can identify conserved immune-organized spatial biomarkers in obstructive lung disease, and that the abundance of these transferred biomarkers scales with independent clinical measures of disease severity.

## 3 Discussion

Here, we present spHOT, a framework for discovering phenotype-associated spatial biomarkers from single-cell resolution spatial transcriptomics (ST). spHOT addresses a key challenge in cohort-scale spatial analysis: disease-relevant tissue structures often occur at heterogeneous and unknown spatial scales, while their cell-or niche-level annotations are often unavailable. To overcome this, spHOT integrates spatial foundation model embeddings, a hierarchical domain tree, and a dual-branch teacher-student multiple instance learning architecture to convert sample-level phenotype labels into spatially coherent importance scores. Across simulations, real tissue benchmarks, and ablation studies, spHOT consistently outperformed existing spatial and single-cell methods in localizing spatial biomarkers. Importantly, it also recovered biologically meaningful tissue organization reported in prior research, including an airway accumulation of *SPP1* + profibrotic macrophages in idiopathic pulmonary fibrosis and ANCA-GN-specific fibrotic-immune interface zones in rapidly progressive glomerulonephritis. To support these findings, we also introduce a metric-based evaluation strategy that quantifies spatial biomarker localization performance rather than relying on qualitative descriptions.

A major strength of spHOT is its label efficiency. While expert annotation is costly and difficult to scale, sample-level labels are readily available in cohort studies. By translating these labels into cell-level importance, spHOT enables quantitative discovery with minimal supervision. This further allows the definition of spatial biomarker burden metrics, such as the proportion of high-scoring cells or the abundance of specific metadomain-defined tissue zones, which can complement conventional histopathology. In addition, spHOT supports cross-disease spatial biomarker transfer. Results within chronic lung diseases suggest that conserved spatial architectures may underlie related disease processes, and further implies that cell scores reflect generalizable tissue organization rather than dataset-specific technical artifacts or cohort-specific characteristics.

Despite these advantages, spHOT has several limitations. It is designed to detect spatially coherent, domain-level cellular communities and is therefore less suited to diffuse signals lacking local spatial organization, where expression-based methods may be more appropriate. Its performance also depends on the representational scope of the underlying spatial foundation model, and generalization to unseen tissue types or platforms remains to be evaluated. Finally, the current spatial autocorrelation metric is applied post hoc for resolution selection. Future work could incorporate spatial coherence directly into the training objective and develop adaptive hierarchies that allocate resolution according to tissue heterogeneity.

Overall, spHOT provides a scalable framework for discovering, comparing, and transferring spatial biomarkers from sample-level labels, addressing a need that grows as ST studies expand to cohort scale.

## 4 Methods

### 4.1 Domain embedding

The spHOT framework starts with a Domain embedding stage that builds an adjacency graph from the spatial transcriptomics (ST) data, extracts spatially-aware cell-level representations, and derives initial domains for subsequent region-level analysis. We utilized Delaunay triangulation with a radius cutoff for the adjacency graph. For each sample *j*, consisting of spatially resolved cells 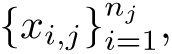 we obtained a 64-dimensional cell-level latent representation *z_i,j_* for each cell *x_i,j_* using a pretrained spatial foundation model.

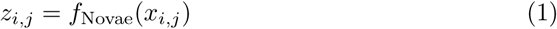

Specifically, we utilized Novae as the latent embedding encoder (v1.0.0; check-point prism-oncology/novae-human-0, Hugging Face revision c35c94b). Novae is a self-supervised foundation model for ST that leverages prototype-based contrastive learning of subgraphs[24]. All embeddings were obtained via zero-shot inference without any task-specific fine-tuning. Novae defines a set of pretrained prototype vectors *P* = *{p*_1_*, p*_2_*,…p*_512_*}*, with each *p_k_ ∈* R^64^, and each cell embedding *z_i,j_* is soft-assigned to these prototypes. The assignment weight between cell *x_i,j_* and prototype *p* is computed as:

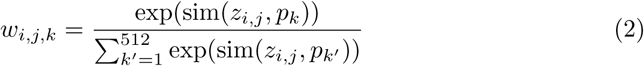

where sim(*·, ·*) denotes cosine similarity. Each cell is then assigned to the initial domain corresponding to the prototype with the highest assignment weight:

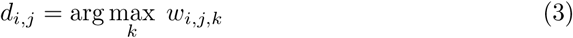

This procedure yields an initial domain set *D* = *{d*_1_*, d*_2_*,…d*_512_*}*. For each initial domain *d ∈ D*, the domain centroid embedding is defined as the mean embedding of all cells assigned to that domain:

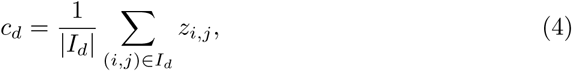

where *I_d_*= *{*(*i, j*) *| d_i,j_*= *d}* denotes the index set of cells assigned to domain *d*. Domains with cells fewer than a minimum threshold were excluded, to ensure robust centroid estimation. For all datasets shown in this work, we set the minimum number of cells as 10 cells, yet no initial domains were excluded (`min cells` in Supplementary Table 1). The resulting domain centroids *{c_d_}_d∈D_* serve as the input for the subsequent Domain tree stage.

### 4.2 Domain tree

To generate metadomains at varying levels of spatial resolution, we performed agglomerative hierarchical clustering on the initial domain centroids obtained from the Domain embedding stage[24]. A distance matrix was constructed using the Euclidean distance between centroids, and Ward’s minimum variance method was applied to iteratively merge the closest clusters, producing a single dendrogram. The Domain tree procedure was implemented using the scipy.cluster.hierarchy module in SciPy[45].

From this dendrogram, sets of metadomains at different levels were derived by cutting the tree at various cluster counts *k ∈ {k*_start_*,…,k*_end_*}* with a fixed step size (step k in Supplementary Table 1). For a given *k*, the dendrogram was partitioned into *k* clusters to define the *k*-level metadomain set *D*^(*k*)^ = *{d_1_*^(*k*)^*, d_2_*^(*k*)^*,…d_k_*^(*k*)^*}*. The centroid embedding of each *k*-level metadomain *d_m_^(k)^* was computed as the mean of the centroid embeddings of its constituent initial domains:

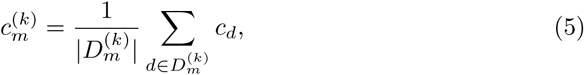

where *d_m_^(k)^* ⊂ D denotes the subset of initial domains merged into *d*^(*k*)^. For each sample, cell-level embeddings were retained, and *k*-level metadomain assignments were used to construct sample-specific metadomain centroids as teacher instances in the subsequent multiple instance learning (MIL) stage. This hierarchical construction enables MIL to be performed at different spatial resolutions.

### 4.3 Multiple instance learning

#### 4.3.1 Problem formulation

We adopted a MIL framework to learn instance-level representations from sample-level binary labels. Each sample *j* can be defined as a bag comprising a set of instances *B_j_* = *{z_i,j_}^nj^*, where *z_i,j_ ∈* R^64^ are the cell-level embeddings obtained during the Domain embedding stage. Each sample was associated with a binary label *y_j_ ∈ {*0, 1*}* (control vs. case), while instance-level labels remained unobserved. All instances were transformed through a shared one-layer MLP encoder *f*_enc_.

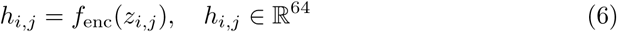

#### 4.3.2 Dual-branch architecture

To simultaneously perform sample-level phenotype prediction and instance-level representation learning, we designed a dual-branch MIL architecture comprising a teacher branch and a student branch that share the encoder *f*_enc_. The two branches are optimized alternately with different objectives: the teacher branch is updated at every training epoch, while the student branch is updated once every three teacher updates (stuOptPeriod in Supplementary Table 1). This alternating optimization ensures that sample-level supervision progressively refines instance-level latent representations through the shared encoder.

#### 4.3.3 Teacher branch

The teacher branch follows the Attention-Based MIL (ABMIL) architecture[33] and operates on metadomain centroids rather than individual cells. For each sample *j*, the sample-metadomain centroid of the *k*-level metadomain *d_m_*^(*k*)^ was defined as the mean embedding of all cells in sample *j* belonging to that metadomain:

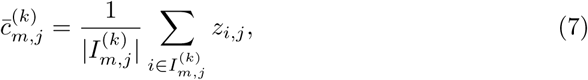

where 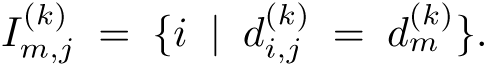 These sample-metadomain centroids were passed through the shared encoder to obtain teacher-branch instances:

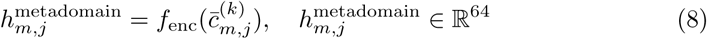

An attention module *f*_attn_ computed normalized attention weights over all metadomain instances within each bag (sample):

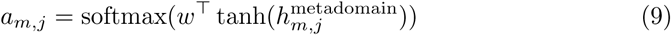

The bag-level representation was obtained as the attention-weighted sum of instance representations, and a classifier *f*_classifier_ produced the sample-level phenotype prediction:

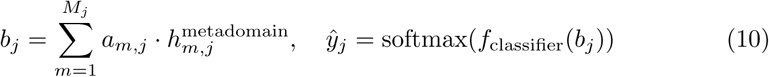

The teacher branch was optimized using the binary cross-entropy loss:

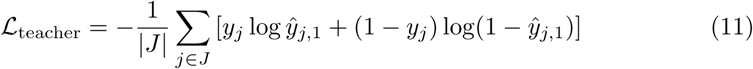

#### 4.3.4 Student branch

The student branch transfers the metadomain-level attention scores learned by the teacher branch to the cell level, enabling instance-level classification. This design follows the philosophy of scMILD[28], aiming to learn representations that effectively separate normal cells, control-like case cells, and case-specific cells in the latent space. Simultaneously, by operating on individual cell embeddings rather than domain centroids alone, it ensures that the shared encoder captures the within-domain cellular variation and avoids overfitting to the domain centroids.

Metadomain-aware cell sampling: The student branch input consists of cell-level embeddings *h*^cell^ = *f*_enc_(*z_i,j_*), constructed via a metadomain-aware cell sampling strategy. For each sample *j*, let *n_d,j_* denote the number of cells in metadomain *d*. The number of cells sampled from metadomain *d* out of a total budget *N*_bag_ was determined as:

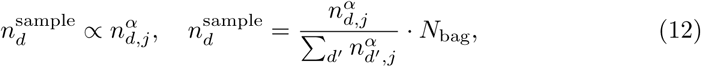

where *α ∈* [0, 1] controls the degree to which the original metadomain proportions are preserved (*α* = 0: uniform allocation; *α* = 1: fully proportional). All *α* values used for results in this work are reported (alpha in Supplementary Table 1). At each epoch, a different random seed was used to sample distinct cell subsets from each domain, providing an augmentation effect that prevents the shared encoder from overfitting to the teacher branch’s metadomain centroids.

Pseudo-labeling and instance classification: Cell-level attention scores *a_i,j_* were extracted from the teacher branch’s attention module and normalized via min-max scaling to yield pseudo instance labels 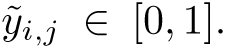 For all instances belonging to negative bags (*y_j_* = 0), pseudo labels were set to zero 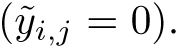 The student branch classifier *f*_student_ produced per-instance predictions 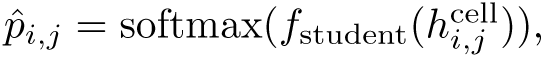 and was optimized using a weighted binary cross-entropy loss *L*_student_.

Orthogonal projection loss (OPL): Within case-labeled bags, cells exhibit varying degrees of contribution, as reflected by their continuous pseudo-labels[46]. To discretize these contributions, a Gaussian Mixture Model (GMM) was fitted to the pseudo-label distribution of case-bag instances, with the number of components selected automatically via the Bayesian Information Criterion (BIC). This procedure naturally distinguishes control-like case cells from case-specific cells. The resulting group assignments, together with negative-bag cells designated as the control group, were used to compute the OPL:

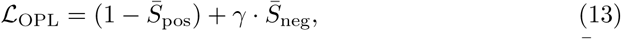

where 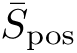 denotes the mean cosine similarity among intra-group instance pairs, 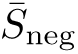denotes the mean absolute cosine similarity among inter-group instance pairs, and *γ* controls the contribution of 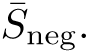 This loss encourages representations within the same group to cluster together while enforcing orthogonality between groups. The total student branch loss was defined as:

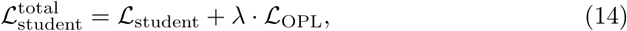

where *λ* controls the contribution of the OPL. In all spHOT runs in this work, *γ* and *λ* values were fixed to 0.5.

#### 4.3.5 Joint optimization

The teacher and student branches were jointly optimized through the shared encoder *f*_enc_ in an alternating fashion. The teacher branch minimized *L*_teacher_ to refine metadomain-level attention scores that identify case-associated spatial biomarkers, while the student branch minimized 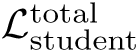 to enforce cell-level separation of control, control-like-case, and case-specific cells. Through this mutual interaction, the teacher’s domain-level attention is progressively transferred to cell-level resolution via the student branch, while the student’s cell-level representation learning reciprocally improves the teacher’s domain-level classification through the shared encoder. Early stopping was applied based on a combined validation metric of bag-level loss and AUROC.

### 4.4 Spatial autocorrelation-based metadomain level selection

To select the best metadomain level (*k^∗^*) from the Domain tree for spatial biomarker discovery, we evaluated each *k* using spatial autocorrelation metrics computed on the cell scores derived from the trained model.

#### 4.4.1 Cell score definition

After training at a given *k*-level, the teacher branch’s attention scores were computed for each sample. For each metadomain *d*^(*k*)^, the attention score *a_m,j_* was min-max normalized and then broadcast to all cells belonging to that metadomain, yielding a cell score *s_i,j_ ∈* [0, 1]. These scores served as input for the spatial autocorrelation metrics.

#### 4.4.2 Moran’s I

Moran’s I is a global spatial autocorrelation statistic that measures the degree to which cell scores of spatially adjacent cells are similar[34]. Given a spatial weight matrix *W* and a cell score vector *s*, Moran’s I is defined as:

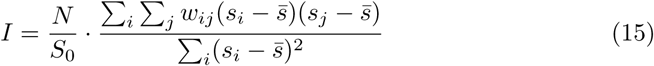

where *N* is the number of cells, *w_ij_* is the spatial weight between cells *i* and *j*, *s̄* is the mean cell score, and 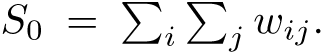 We adjusted the negative values by *I*_norm_ = (*I* + 1)*/*2.

#### 4.4.3 Geary’s C

Geary’s C is a local dissimilarity-based statistic that directly measures the difference in cell scores between neighboring cells[35]:

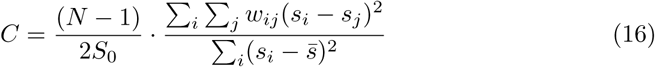

Geary’s C typically ranges from 0 to approximately 2, with values closer to 0 indicating greater spatial similarity among neighboring cells. To limit the transformed values to [0, 1], we first clipped *C* to the interval [0, 2] and then applied the following transformation:

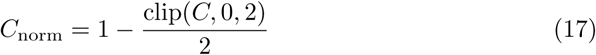

#### 4.4.4 Spatial score and early stopping

For each metadomain level *k*, a composite spatial score was computed as the equally weighted average of the two normalized metrics across the case samples within the validation set:

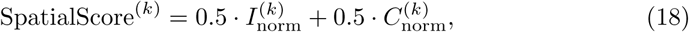

where each metric value represents the median across case samples within each fold, averaged across folds. As a sanity check, a minimum validation AUROC threshold was enforced to exclude granularity levels at which the classification performance was excessively low, prioritizing spatial coherence while filtering out degenerate solutions (auroc threshold in Supplementary Table 1). The final *k^∗^* was selected as the value maximizing the spatial score. An early stopping criterion was applied: if no improvement in the spatial score was observed for *P_k_* consecutive values of *k*, the search was terminated to avoid unnecessary computation (patience k in Supplementary Table 1).

### 4.5 Simulation framework

#### 4.5.1 Expression generation

For all simulation dataset generation, we employed the scCube Python package (v2.0.0)[36]. Simulated gene expression profiles covering 974 genes were generated using scCube trained on a CosMx-profiled inflammatory bowel disease (IBD) pouch dataset[37], from which seven cell types were selected and anonymized as types A through G. To minimize sample-level batch effects, expression was generated in a single pooled step: the total cell count of each cell type across all samples was summed, a single large sample matching this total count and per-cell-type count was generated through scCube gene expression generation module. Subsequently, the resulting expression vectors were randomly assigned back to individual samples according to each cell’s type label. Each sample contained between 2,000 and 4,000 cells, with the exact count drawn randomly per sample.

#### 4.5.2 Spatial layout construction

Each simulated sample consists of cells placed within a 100 *×* 100 pixel field. Control samples were simulated to contain all seven cell types randomly distributed at proportions matching the source dataset. Case samples were constructed identically, except that 50% of each of two selected cell types were repositioned into defined spatial structures constituting the ground truth spatial biomarkers. Three structure morphologies were created using different shape options and functions of scCube. Circular aggregates (Circle option in generate pattern custom cluster), vessel-like elongated formations (generate pattern custom stripes), and random mixing of cell types (generate pattern custom mixing). In the spatial-only circular scenario, 1–4 circular aggregates were placed per sample with total biomarker area held constant. This resulted in more numerous circles becoming proportionally smaller. In the mixed circular and vessel-like scenario, a subset of circles were replaced by vessel-like elongated structures of equivalent cell count: one circle was replaced in the two-circle setting, one or two in the three-circle setting, and two in the four-circle setting. In the vessel-only scenario, all structures were created vessel-like. Spatial biomarkers were constrained to be non-overlapping.

#### 4.5.3 Confounding scenarios

Two additional scenarios progressively introduce non-spatial discriminative signals. In the circular and composition altered scenario (CC), spatial biomarkers were created to follow the circular-only design, but the proportion of the two constituent cell types was increased in case samples relative to controls, introducing a compositional difference detectable without spatial information. In the circular and composition altered with perturbation scenario (CCP), one cell type (F) in case samples was additionally replaced by a stressed variant (F*) expressing elevated levels of heat shock protein genes *HSPA1A/B*, introducing a transcriptional perturbation on top of the compositional shift.

### 4.6 Benchmarking analysis

Seven types of existing methods were benchmarked alongside spHOT, evaluating the performance of spatial biomarker detection. For sample-level classification benchmarking, two additional methods were included (ProtoCell4P, Hier-MIL). Identical sample fold assignments were used across all methods. HiDDEN and STANDS were excluded from fold assignment as they are self-supervised methods without a sample-level classifier. For all datasets, each method was run using the default hyperparameters specified in its respective tutorial, unless otherwise stated. Below are descriptions specific to each method.

SPACE-GM[29] was run using the authors’ GitHub implementation, cloned at commit 9d1132f6. We used two different graph node feature settings, cell type labels (CT) and gene expression (GEX), treating them as separate comparisons. The microenvironment prediction values produced by SPACE-GM on held-out test case samples were used as cell importance scores, as previously shown in the original paper.

STANDS[30] (v1.1.0) is implemented as a Python package. STANDS was trained on control samples only using its GAN-based self-supervised framework, after which case samples were evaluated to identify anomalous cells producing large reconstruction errors. These STANDS anomaly scores in the case samples were used as cell importance scores.

scMILD[28] was evaluated using the authors’ implementation at git commit 127971c. scMILD attention scores from its MIL framework were used directly as cell importance scores.

We employed HiDDEN[25] from the authors’ implementation at git commit 326a1ef. HiDDEN was run with three embedding strategies (PCA, linear scVI, and scVI), using the scvi-tools[47] Python package (v1.3.3). Logistic regression option was used as a cell-level classifier. Per-cell continuous perturbation scores were used as cell importance scores for evaluation.

For sample classification performance evaluation, existing single-cell-based sample classification methods, Hier-MIL[26] (git commit 2fb1558) and Protocell4P[38] (git commit 1094489), were added for benchmarking. ProtoCell4P was run with the number of prototypes selected from 4, 8, 16, 50 to approximately match the number of annotated cell types per dataset, following ProtoCell4P’s design intent of capturing one characteristic cellular state per prototype.

### 4.7 Evaluation metrics

Cell score quality was assessed using three primary metrics and two supplementary metrics, applied to each test-set case sample independently. For each metric, the best-performing method was assigned to that of best median performance. ES, AUPRC, and median ground truth percentile are reported in main figures. NES and AUROC are reported in Extended Data Fig. 3 and Extended Data Fig. 4.

#### 4.7.1 Enrichment score (ES)

Cells are ranked by predicted score in descending order. A GSEA[48]-style running sum is computed by incrementing by 1*/n*_pos_ for each ground truth spatial biomarker cell encountered and decrementing by 1*/n*_neg_ for each non-biomarker cell, where *n* denotes the number of cells per sample. ES is defined as the maximum of this running sum, capturing whether biomarker cells are concentrated among the highest-scoring cells.

#### 4.7.2 Normalized enrichment score (NES)

The Normalized enrichment score (NES) is obtained by comparing the observed ES against a permutation null distribution (1,000 permutations of ground truth labels with score rankings held fixed), reported as a z-score: NES = (ES *− µ*_null_)*/σ*_null_. Permutation *p*-values are computed as (*k* + 1)*/*(*n*_perm_ + 1), where *k* is the number of permuted ES values equal or greater than observed ES.

#### 4.7.3 Area under the precision-recall curve (AUPRC)

Precision-recall curves are computed over continuous cell scores with ground truth biomarker membership as the binary label. AUPRC provides a stricter assessment than ES: a method that assigns uniformly high scores to all cells would achieve high ES but low AUPRC, as false positives are explicitly penalized. The random baseline equals the positive prevalence in each sample[49].

#### 4.7.4 Median ground truth percentile

Each cell is assigned a percentile rank based on its cell score (rank0 = highest score). The median percentile across all ground truth spatial biomarker cells is reported, where lower values indicate better prioritization.

#### 4.7.5 AUROC

The standard area under the receiver operating characteristic curve is computed over continuous cell scores against ground truth labels of spatial biomarker cells[50].

### 4.8 Cross-validation and statistical testing

Each simulation scenario used five-fold cross-validation with 12 training, 4 validation, and 4 test samples per condition per fold. In the Xenium IPF dataset, 10 control samples and 16 case samples were split to five-folds, regardless of disease severity. Multiple samples originating from the same patient were grouped to the same fold. In the CosMx DKD dataset, 3 control samples and 14 case samples were split to three-folds, regardless of DKD class. In the Xenium RPGN dataset, 6 healthy controls, 19 SLE-GN, and 32 ANCA-GN samples were split to three-folds. In both comparison settings, the ANCA-GN (case) samples were fixed in each fold, only replacing the control samples. For all datasets, each case sample appeared in the test set of exactly one fold, yielding non-overlapping per-sample evaluations per box plot. Pairwise method comparisons were performed using the Wilcoxon signed-rank test (paired, two-sided).

### 4.9 Ablation studies

Three ablation studies were designed to evaluate the individual contributions of spHOT’s core architectural components. Each ablation was applied to both the simulation dataset (circular, mixed circular and vessel-like scenarios) and the Xenium IPF dataset, using identical cross-validation folds and evaluation metrics as the main benchmarking experiments.

Ablation 1 (without domain tree): The hierarchical domain tree was removed and cell-level MIL was applied directly to Novae cell embeddings. This setting is similar to scMILD, but uses pretrained foundation embeddings instead of pure autoencoder based latent features. The teacher branch operated on cell embeddings rather than metadomain centroids, and no metadomain-level spatial abstraction was performed.

Ablation 2 (without metadomain-aware cell sampling): The domain tree and dual-branch architecture were retained, but metadomain-aware cell sampling was removed from the student branch. All cells within each sample were used for student training in every epoch.

Ablation 3 (without student branch): The student branch was removed entirely. Sample-level classification was performed using attention-based MIL on metadomain centroids via the teacher branch only. Cell-level scores were derived from the teacher’s metadomain-level attention weights assigned to each cell’s parent metadomain, without cell-level representation refinement or OPL-based regularization.

Training time per epoch was recorded separately for teacher and student branches across all settings. Cell score quality was evaluated using the same three primary metrics (ES, AUPRC, median ground truth percentile) on identical test-set samples. All experiments were conducted on a Linux server equipped with 2 AMD EPYC 7763 CPUs (64 cores each), NVIDIA RTX A6000 GPUs with 48 GB memory, and approximately 1 TB of system memory.

### 4.10 Public dataset collection and preparation

#### 4.10.1 CosMx inflammatory bowel disease (IBD) pouch dataset

We utilized the CosMx IBD pouch dataset from Olivas et al.[37]. This public dataset was used to generate gene expression profiles and cell type labels in the simulation dataset. In order to generate robust and reliable gene expression space, we filtered out cells expressing less than 50 transcripts, resulting in total 308,091 cells. No gene filtering was conducted and all 974 genes were retained. Cell type labels were preserved from the original study.

#### 4.10.2 Xenium idiopathic pulmonary fibrosis (IPF) dataset

The Xenium IPF dataset was obtained from Vannan et al.[39], comprising 10 control lung samples and 16 IPF lung samples profiled using the 10x Genomics Xenium platform with a 343-gene panel, totaling 823,728 cells. All gene panels were utilized without gene filtering, and cell type annotations in the original dataset were directly used. Ground truth immune hotspot regions were derived from clinician-performed histopathological annotations provided in the original study’s cell-level metadata. TLS annotations were available in 2 of the 16 IPF samples, while mixed inflammation annotations were available in 9 out of 16 IPF samples. Only one sample possessed both TLS and mixed inflammation.

#### 4.10.3 CosMx diabetic kidney disease (DKD) dataset

The CosMx DKD dataset was obtained from Meadows et al.[42], comprising 3 control kidney samples and 14 DKD samples profiled using the NanoString CosMx Spatial Molecular Imager. Total 1008-gene panel was profiled in 358,706 healthy and DKD kidney cells. The original study’s cell type annotations were preserved and directly used. No gene filtering was applied. Ground truth tubular injury regions were constructed following the spatial neighborhood framework established by Meadows et al., who identified cells adjacent to *HAVCR1+* injured proximal tubular (iPT) cells within a 30*µm* radius to characterize the injury microenvironment. Using the same approach, iPT cells from the original study’s cell-level annotations were identified and their 30*µm* neighborhoods were computed via KDTree-based spatial queries, using the scipy.spatial[45] Python package (v1.15.3). All cells within these neighborhoods were labeled as positive for tubular injury, with remaining cells as negative (background).

#### 4.10.4 Xenium rapidly progressive glomerulonephritis (RPGN) dataset

We utilized 6 healthy controls, 19 SLE-GN, and 32 ANCA-GN kidney samples from the Xenium RPGN dataset, obtained from Sultana et al.[43]. This dataset profiled kidney tissues using the 10x Genomics Xenium platform with a 480-gene panel, totaling 2,755,475 cells. We directly utilized the cell type and niche annotations reported in the original study. No gene filtering was applied.

#### 4.10.5 Xenium chronic obstructive pulmonary disease (COPD) dataset

The Xenium COPD dataset was obtained from Zhang et al.[44], comprising four tissue microarray slides containing 99 cores from 38 patients and donors spanning GOLD stages 0 to IV, profiled using the 10x Genomics Xenium platform with a 480-gene panel. We reconstructed the full analysis from the raw outputs, with only the core-level GOLD stage and emphysema category known from the original study. Cells were assigned to cores using rectangular spatial bounding boxes defined per slide from the cell centroid coordinates, yielding all 99 cores. Quality control followed the thresholds reported in reference: transcripts were retained at Phred *Q >* 20, and cells were required to have more than 5 detected features, more than 9 total counts, and a nuclear area between 6 and 80*µm*^2^. All gene panels were utilized. Filtered count matrices were log-normalized, scaled, and reduced by principal component analysis (50 components). Using the Scanpy[51] Python package (v1.11.5), the neighbor graph was constructed and embedded with UMAP, and cells were clustered using the Leiden algorithm (resolution = 0.5). Canonical marker genes from the original study were examined across Leiden clusters to assign major lineage identities. To resolve finer cell states, each major cluster was sub-clustered by re-running Leiden clustering within the initial cluster, and the resulting subclusters were annotated by marker expression, including inflammatory nonimmune states identified by elevated NF-κB, interferon, and cytokine program genes. The final processed dataset comprised 816,245 cells annotated into 34 cell types, including canonical lineages, disease-associated states, and inflammatory subtypes (Supplementary Fig. 1–2).

### 4.11 Metadomain tree lineage analysis

To trace the emergence of spatial biomarkers across metadomain levels, individual metadomain lineages were tracked through the hierarchical domain tree. Starting from a selected metadomain at the finest resolution (final level for the MIL run), parent metadomains at each coarser resolution were identified via the Ward’s linkage matrix. For each metadomain in a lineage, cell-level scores were averaged across case samples within the fold to generate a single tree (Fig. 4a). Tree was visuzlized using GraphViz[52]. Spatial visualizations were generated at selected *k* levels to illustrate the progressive resolution of the macrophage niche, with cells belonging to the selected metadomain lineage highlighted and all other cells displayed in gray (Fig. 4b,c).

#### 4.11.1 Cell label proportion projection on domain tree

For each metadomain at each resolution *k*, the proportion of each annotated cell (cell type or cell-level clinician annotation) was computed and projected onto the corresponding tree node. This enabled visual assessment of whether high-scoring metadomain lineages corresponded to biologically coherent cell type enrichments.

#### 4.11.2 Level-dependent enrichment quantification

To quantify the level dependence of macrophage niche detection, the ES metric was computed at each *k* level for each case sample, using the cell scores of cells within the target metadomain. Per-sample ES values were averaged across case samples and plotted as a function of *k* for visualization (Fig. 4f).

### 4.12 Metadomain-level high/low classification

Within each fold, a two-component Gaussian mixture model was fitted to the metadomain-level mean scores, and each metadomain was classified as high or low based on its posterior assignment. Final metadomain classifications were determined by majority voting across all folds: a metadomain was assigned as high if it was classified as high in two or more out of three folds, and low otherwise. This procedure was applied identically to the CosMx DKD dataset (three-folds, 17 metadomains) and the Xenium RPGN dataset S2 (three-folds, 25 metadomains).

### 4.13 Multi-context comparison

Two independent spHOT experiments were run on the 32 ANCA-GN samples with different control groups. Setting 1 (S1) used 6 healthy controls (Healthy vs. ANCA), while Setting 2 (S2) used 19 SLE-GN samples (SLE vs. ANCA). Each setting was run with the full spHOT pipeline with hyperparameters reported in Supplementary Table 1. The selected metadomain resolutions were *k* = 19 for S1 and *k* = 25 for S2.

### 4.14 Pseudobulk differential expression and pathway enrichment analysis

Raw counts of cells were aggregated per sample to create pseudobulk expression profiles. In the Xenium RPGN dataset, the glomerular and periglomerular cells were selected and aggregated per sample. Differential expression analysis was performed using the PyDESeq2[53] Python package (v0.5.4). Genes were ranked according to the differential expression statistic and used for preranked gene set enrichment analysis with the GSEApy[54] Python package (v1.1.12). Enrichment analysis was performed against the Reactome[55] (v2026.1) and Gene Ontology Biological Process[56][57] (GO-BP) (v2026.1) gene set database downloaded from MSigDB[48][58]. Gene sets with FDR *<* 0.05 were considered significant. The enrichment results were visualized as a barplot of normalized enrichment scores (NES) for the top-ranked pathways.

### 4.15 Metadomain-level expression analysis

Expression of gene sets was visualized across the metadomains with a dot plot, where dot size represents the fraction of cells expressing each gene and color intensity represents mean expression. For analysis we used the Scanpy[51] Python package (v1.11.5). This analysis was used to identify active/chronic injury signatures in CosMx DKD dataset, and concentrated ECM gene expression and adaptive immune system in the Xenium RPGN dataset.

### 4.16 Zone identification and neighborhood enrichment analysis

In the Xenium RPGN dataset, functional zone categories were defined by integrating metadomain-level cell-type composition, pathway activity, and spatial neighborhood enrichment. For each metadomain, we quantified the proportions of annotated cell types, and summarized pathway gene expression at the metadomain level. Spatial neighborhood enrichment between metadomain pairs was computed across 32 ANCA-GN samples using squidpy.gr.spatial neighbors from the Squidpy[59] Python package (v1.8.1). For each sample, observed frequencies of spatial adjacency between metadomain pairs were compared with random expectation and reported as z-scores. Sample-wise z-scores were then averaged across samples and visualized as a heatmap.

Metadomains were assigned to three functional zone categories. Fibrotic core metadomains were defined by enrichment of fibrotic mesangial cells and fibroblasts together with high ECM pathway activity. Immune hotspot metadomains were defined by high adaptive immune system pathway activity with enrichment of macrophages, T cells and B cells. Fibrotic-immune interface metadomains were defined as metadomains spatially adjacent to both fibrotic core and immune hotspot metadomains, with elevated ECM/adaptive immune system pathway activity but without strong enrichment of fibrotic mesangial cells in their cell type composition.

### 4.17 Cross-disease application of spHOT

To assess cross-disease generalizability, the spHOT teacher branch trained on the Xenium IPF dataset was applied to the Xenium COPD dataset. We used the teacher branch from the resolution and fold that resolved the profibrotic macrophage niche in IPF (*k* = 21), ensuring that the transferred model was identical to the one characterized in the IPF analysis. Zero-shot inference was performed for all COPD at the single-cell level, using the same configuration applied to the IPF data (Supplementary Table 1). For each core, the cell embeddings were passed through the trained encoder and the teacher attention module to obtain a per-cell score, without aggregation to metadomains. Scores were min-max normalized across all cells, and a two-component Gaussian mixture model was fitted to the normalized scores to binarize cells into high and low groups, with the higher-mean component designated as the high group. Per-core high cell burden was defined as the fraction of high cells among all cells within a core. Cell type composition of high versus low cells was computed per core and compared by two-sided Mann-Whitney U tests on per-core proportions with Benjamini-Hochberg correction across cell types. For severity association, per-core high cell burden was correlated with GOLD stage and emphysema category using Spearman correlation, and dichotomized comparisons (GOLD 0/I/II versus III/IV; emphysema 0/1 versus 2/3) were assessed by two-sided Mann-Whitney U tests.

## 5 Data availability

The datasets analyzed in this study were mainly obtained from previously published sources. The CosMx inflammatory bowel disease (IBD) pouch dataset[37] is available from the Gene Expression Omnibus (GEO) under accession GSE283625. The Xenium idiopathic pulmonary fibrosis (IPF) dataset[39] is available from GEO under accession GSE250346. The CosMx diabetic kidney disease (DKD) dataset[42] is available from GEO under accession GSE325587. The Xenium rapidly progressive glomerulonephritis (RPGN) dataset[43] is available from GEO under accession GSE294965. The Xenium chronic obstructive pulmonary disease (COPD) dataset[44] is available from GEO under accession GSE313006. All simulation datasets generated in this study, including the frozen source expression profiles and simulated spatial layouts, are deposited at Zenodo[60]. Gene set annotations used for pathway enrichment analysis were obtained from MSigDB[58](Reactome and GO Biological Process collections).

## 6 Code availability

spHOT is implemented in Python and can be publicly accessed in the Github repository (https://github.com/DHKim327/spHOT). All tutorial notebooks for running spHOT and reproducing figures can be found in GitHub.

## 7 Acknowledgements

This work was supported by the National Research Foundation of Korea (NRF) grant funded by the Korea government (MSIT) (RS-2026-25473777 and RS-2026-25516068), and by a grant from the Korea Health Technology R&D Project through the Korea Health Industry Development Institute (KHIDI), funded by the Ministry of Health & Welfare, Republic of Korea (RS-2024-00438641). The authors thank all members of the lab, especially Kyeonghun Jeong and Ari Hong, for their helpful discussions and feedbacks during the development and writing process of this work.

---

**Extended Data Fig. 1:**
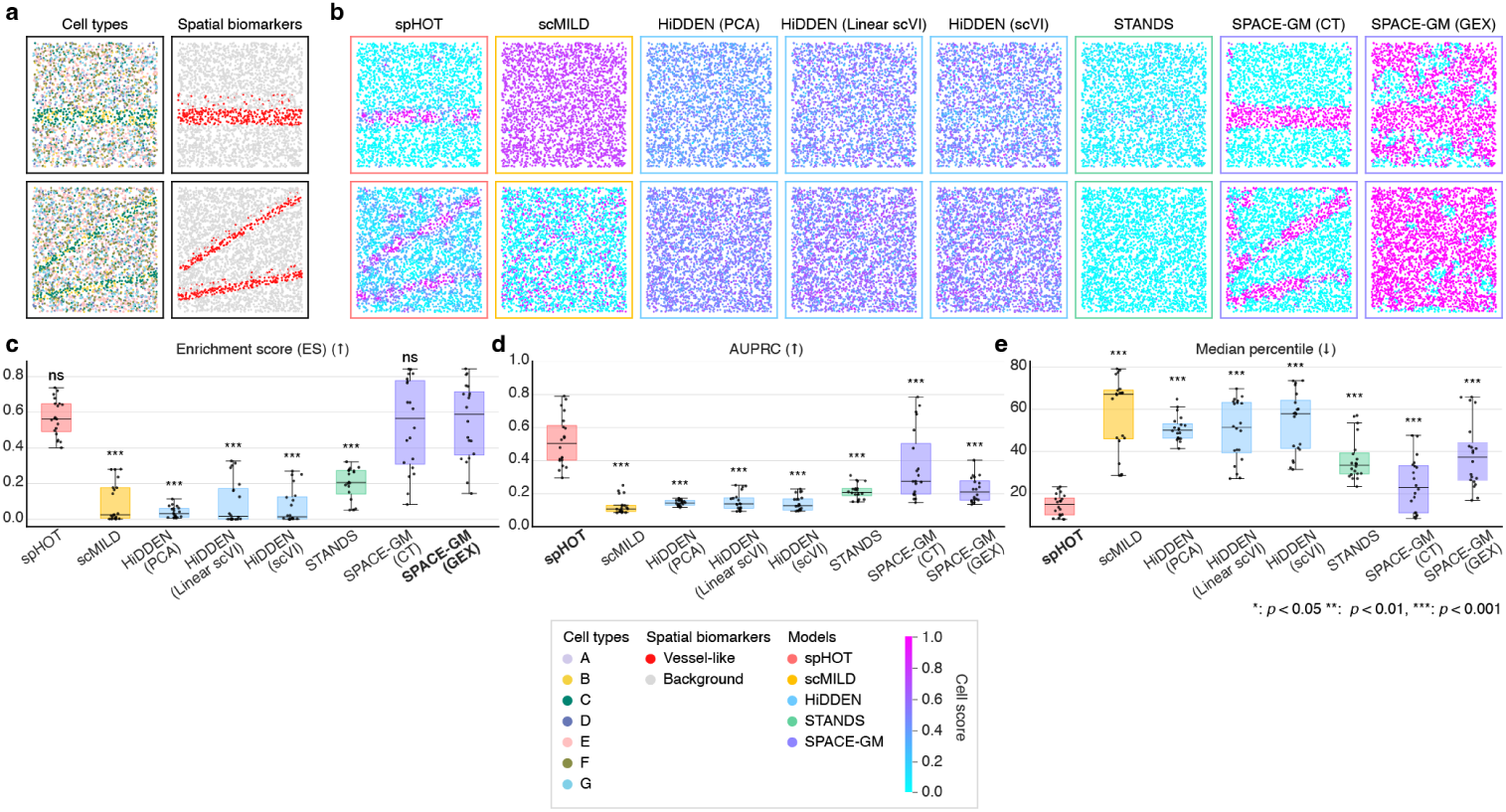
Performance evaluation of vessel-like spatial biomarker discovery on simulated datasets. **a** Two representative spatial lay-outs, colored by cell type (left) and ground truth spatial biomarker (right). **b** Continuous heatmaps of normalized cell scores within [0, 1] range for all methods. Box outlines indicate method type. Upper and lower rows correspond to the same representative samples as in **a**. **c**–**e** Benchmarking performance across 20 test-set case samples, showing **c** ES; **d** AUPRC; **e** median ground truth percentile. Box plots show the median (centerline), interquartile range (box), and 1.5*×* IQR (whiskers); each point is one sample. The best-performing method is highlighted in bold. Asterisks denote significant differences from the best-performing method (paired Wilcoxon signed-rank test; \**p <* 0.05, \*\**p <* 0.01, \*\*\**p <* 0.001, ns, not significant).

**Extended Data Fig. 2:**
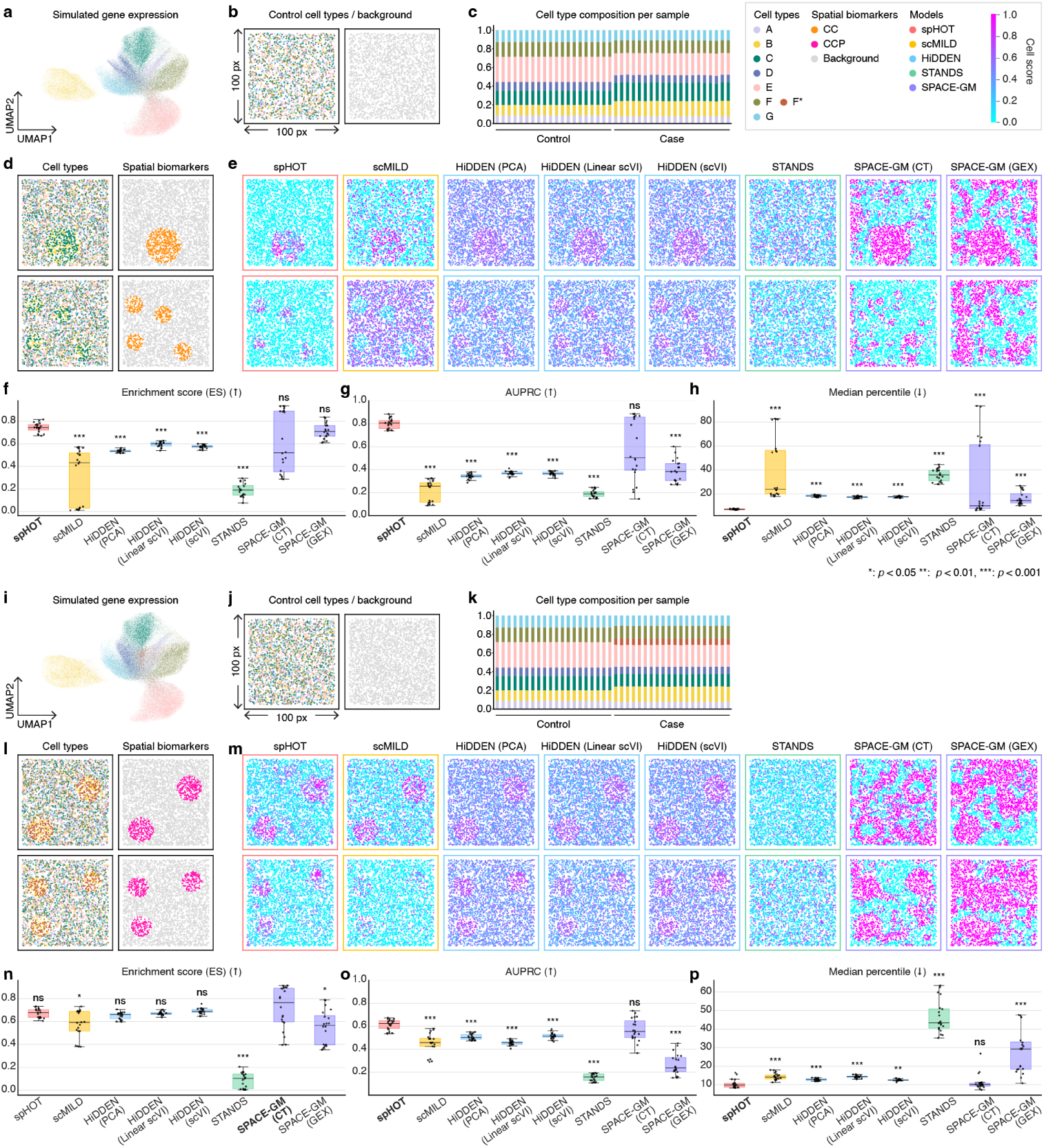
Performance evaluation of spatial biomarker discovery on simulated datasets with confounding scenarios. **a**, **i** UMAP visualization of simulated gene expression profiles for CC (**a**) and CCP (**i**) generated using scCube, comprising seven anonymized cell types (A–G) and one perturbed cell type (F*). **b**, **j** Representative spatial layouts of a control sample for CC (**b**) and CCP (**j**), colored by cell type (left) and ground truth spatial biomarker (right), showing a random distribution of all cell types. **c**, **k** Cell type composition per sample across all control and case samples for CC (**c**) and CCP (**k**). Note cell type composition shift observed in case samples. **d**–**h**, **l**–**p** Evaluation on the CC spatial biomarker scenario (**d**–**h**) and CCP scenario (**l**–**p**). **d**, **l** Two representative spatial layouts, colored by cell type (left) and ground truth spatial biomarker (right) for CC (**d**) and CCP (**l**). **e**, **m** Continuous heatmaps of normalized cell scores within [0, 1] range for all methods for CC (**e**) and CCP (**m**). Box outlines indicate method type. Upper and lower rows correspond to the same representative samples as in **d** and **l**. **f** –**h**, **n**–**p** Benchmarking performance across 20 test-set case samples, showing **f**3,8**n** ES; **g**, **o** AUPRC; **h**, **p** median ground truth percentile. Box plots show the median (centerline), interquartile range (box), and 1.5*×* IQR (whiskers); each point is one sample. The best-performing method is highlighted in bold. Asterisks denote significant differences from the best-performing method (paired Wilcoxon signed-rank test; \**p <* 0.05, \*\**p <* 0.01, \*\*\**p <* 0.001, ns, not significant).

**Extended Data Fig. 3:**
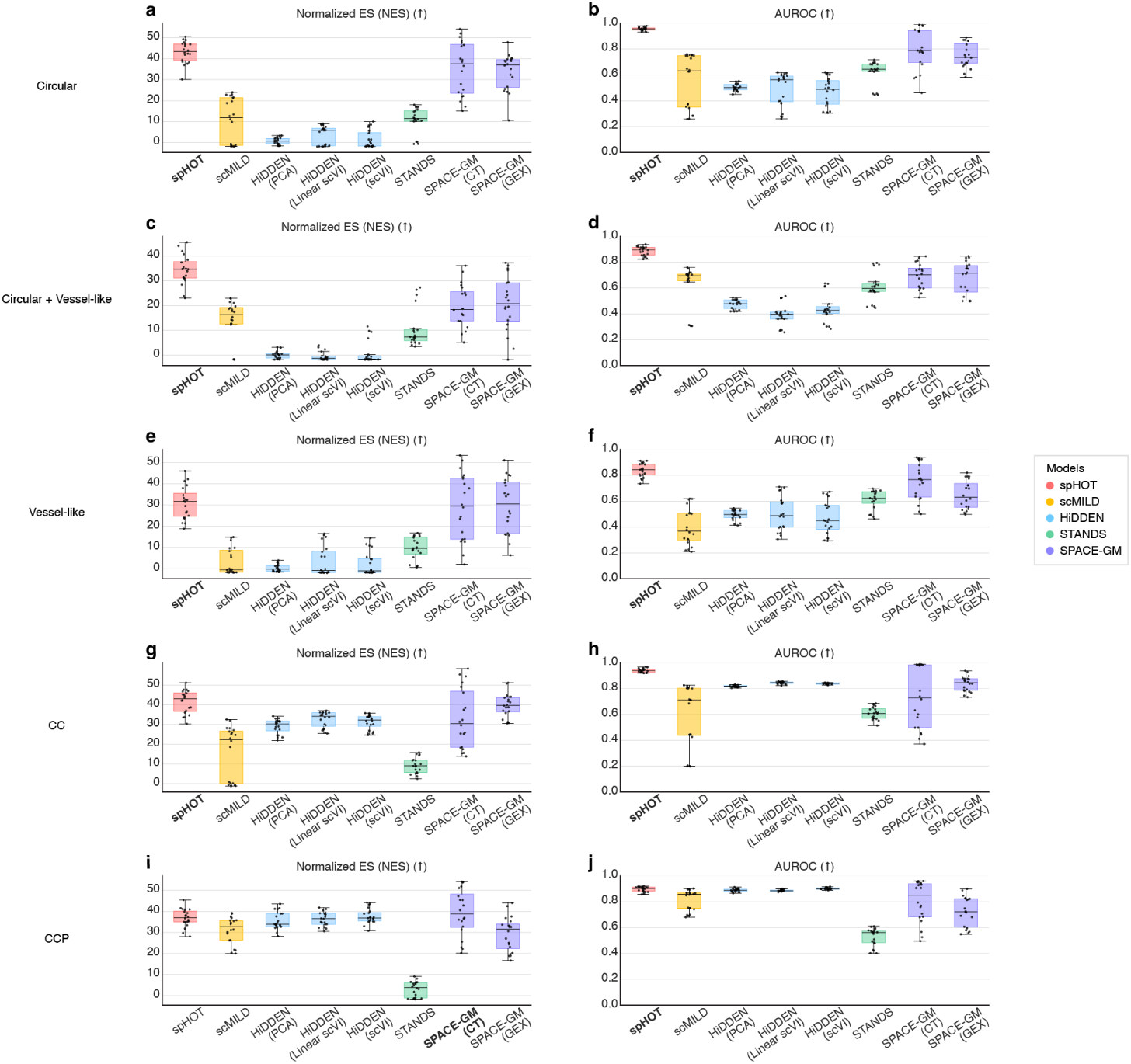
Performance evaluation of spatial biomarker discovery on all simulated datasets with different metrics. **a**–**b** Benchmarking performance of circular-only scenario across 20 test-set case samples, showing **a** Normalized enrichment score (NES); **b** Area under the receiver operating characteristic curve (AUROC). **c**–**d** Benchmarking performance of mixed circular and vessel-like scenario across 20 test-set case samples, showing **c** NES; **d** AUROC. **e**–**f** Benchmarking performance of vessel-like scenario across 20 test-set case samples, showing **e** NES; **f** AUROC. **g**–**h** Benchmarking performance of CC spatial biomarker scenario across 20 test-set case samples, showing **g** NES; **h** AUROC. **i**–**j** Benchmarking performance of CCP spatial biomarker scenario across 20 test-set case samples, showing **i** NES; **j** AUROC. The best-performing method is highlighted in bold. Box plots show the median (centerline), interquartile range (box), and 1.5*×* IQR (whiskers); each point is one sample.

**Extended Data Fig. 4:**
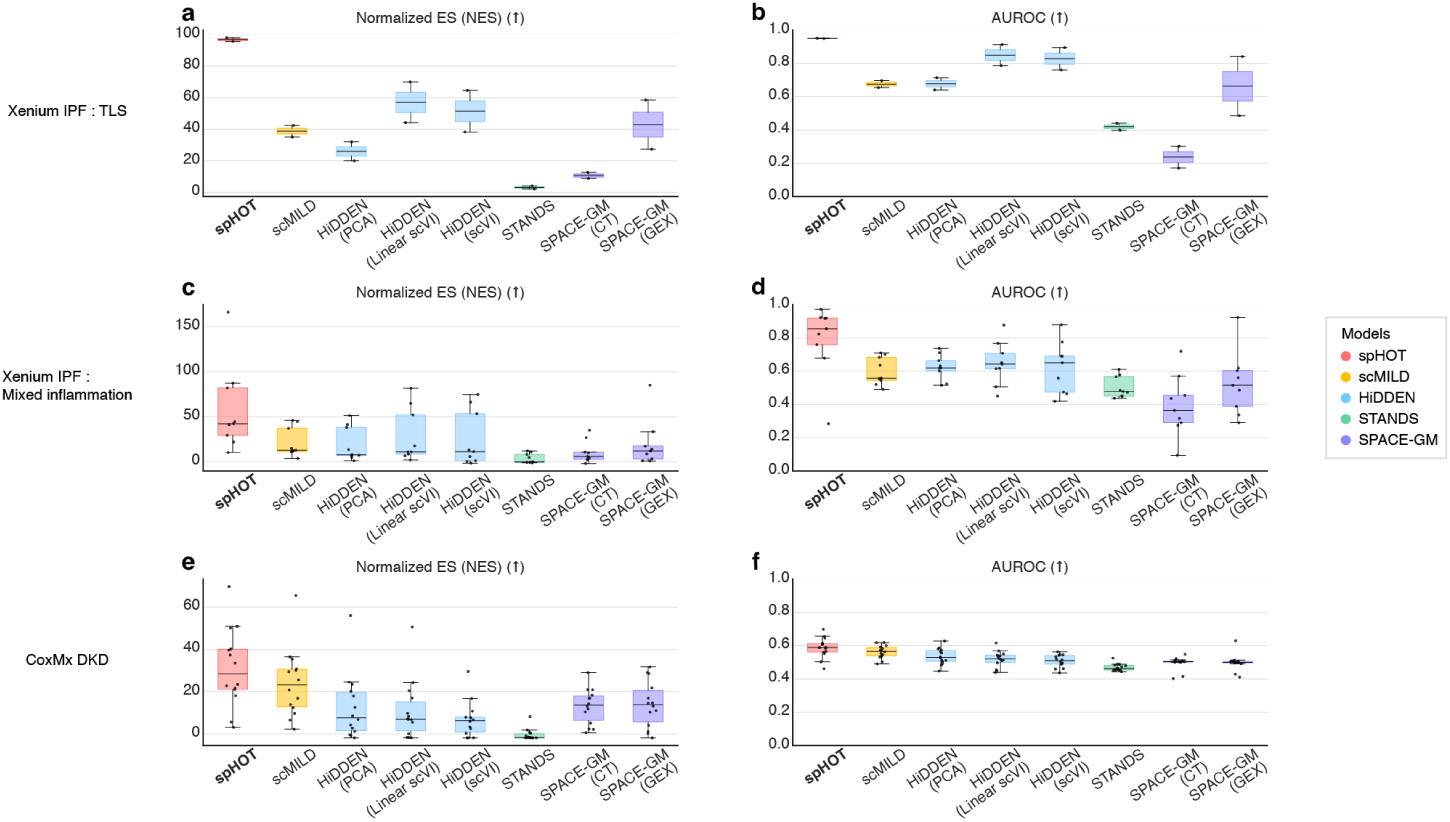
Performance evaluation of spatial biomarker discovery on all real world datasets with different metrics. **a**–**b** Benchmarking performance of Xenium IPF dataset TLS ground truth discovery across 2 test-set case samples, showing **a** Normalized enrichment score (NES); **b** Area under the receiver operating characteristic curve (AUROC). **c**–**d** Benchmarking performance of Xenium IPF dataset mixed inflammation ground truth discovery across 9 test-set case samples, showing **c** NES; **d** AUROC. **e**–**f** Benchmarking performance of CosMx DKD dataset tubular injury ground truth discovery across 14 test-set case samples, showing **e** NES; **f** AUROC. The best-performing method is highlighted in bold. Box plots show the median (centerline), interquartile range (box), and 1.5*×* IQR (whiskers); each point is one sample.

**Extended Data Fig. 5:**
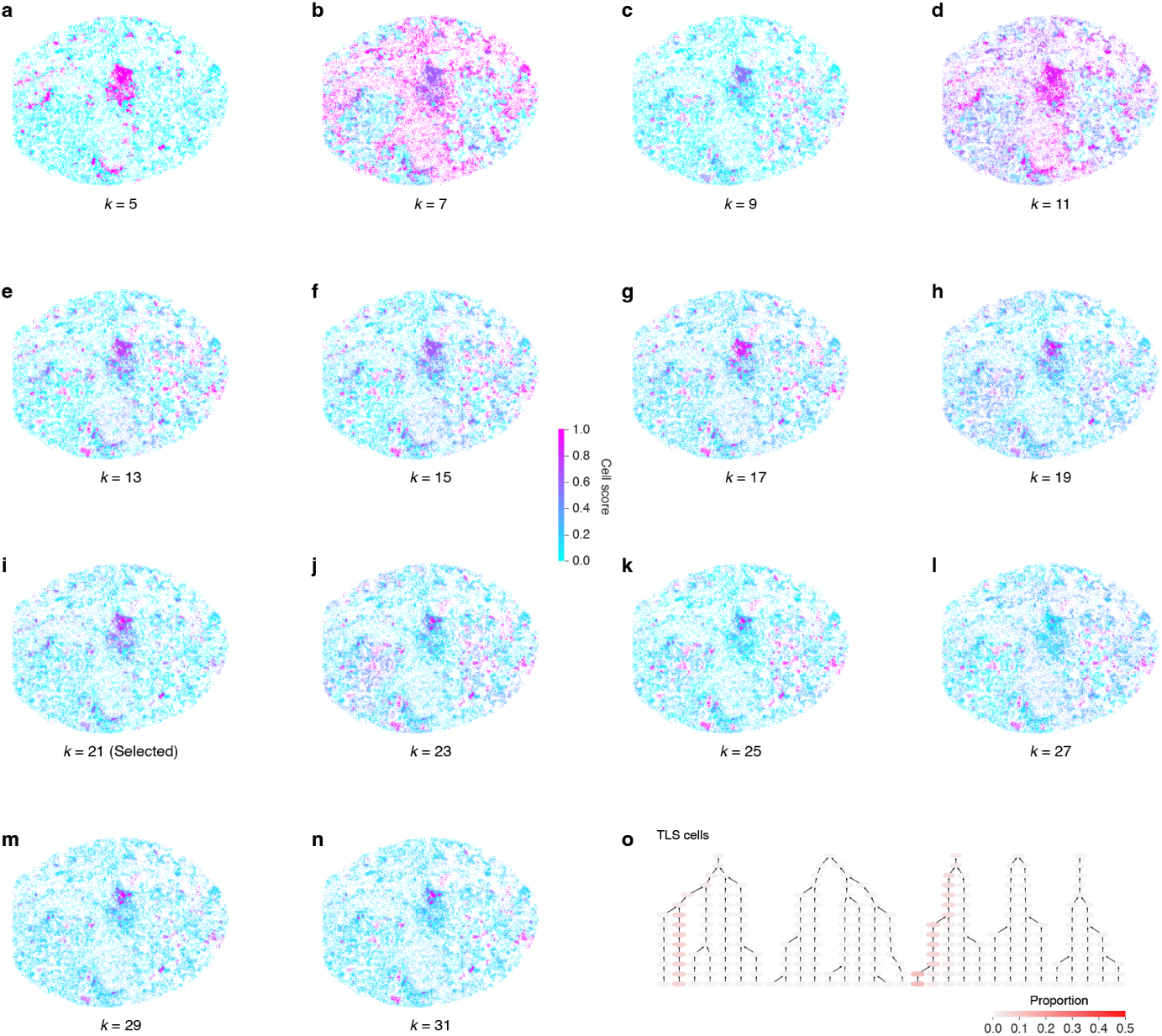
Total cell score maps across successive metadomain resolutions. **a**–**n** Cell score spatial visualizations at successive metadomain resolutions (*k* = 5 (**a**); 7 (**b**); 9 (**c**); 11 (**d**); 13 (**e**); 15 (**f**); 17 (**g**); 19 (**h**); 21 (**i**); 23 (**j**); 25 (**k**); 27 (**l**); 29 (**m**); 31 (**n**)) for representative sample VUILD91MA. Scores are normalized to [0, 1] and displayed as a continuous heatmap. **o** TLS-annotated cell proportion projections onto the metadomain tree.

**Extended Data Fig. 6:**
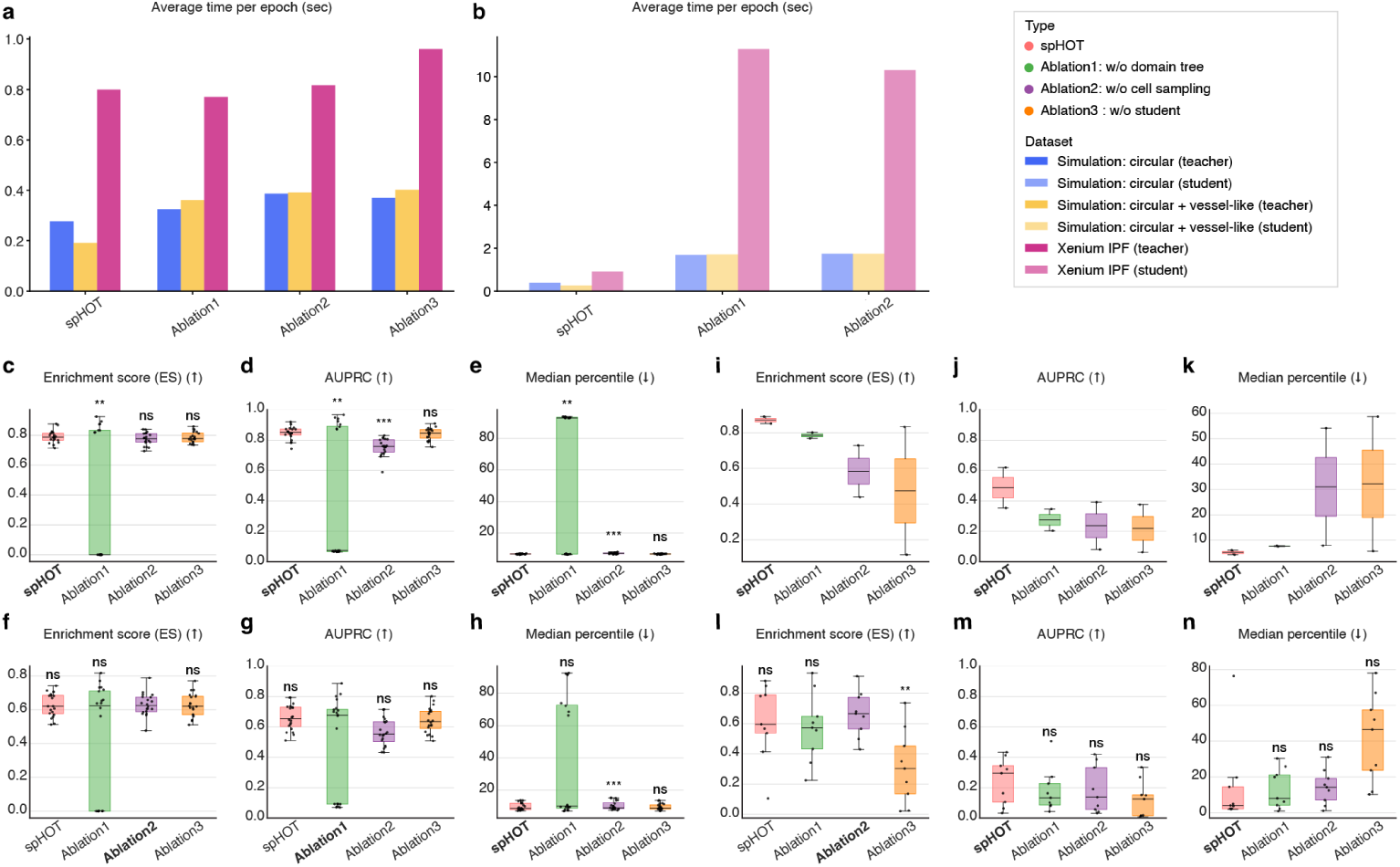
Ablation studies evaluating the contribution of spHOT key architectural components. **a**,**b** Average training time (seconds) per epoch for spHOT and three ablation settings on the simulation dataset (circular and mixed circular and vessel-like scenarios) and Xenium IPF dataset, summarized by (**a**) teacher branch and (**b**) student branch separately. Ablation 3 (without student branch) has no student branch training time. **c**–**e** Cell score performance of spHOT and ablation settings on the circular-only scenario simulation across 20 test-set case samples, showing **c** ES; **d** AUPRC; **e** median ground truth percentile. **f** –**h** Cell score performance on the mixed circular and vessel-like scenario across 20 test-set case samples, formatted as in **c**–**e**. **i**–**k** Cell score performance on the Xenium IPF TLS ground truth discovery across 2 test-set case samples, formatted as in **c**–**e**. **l**–**n** Cell score performance on the Xenium IPF mixed inflammation ground truth discovery across 9 test-set case samples, formatted as in **c**–**e**. Box plots show the median (center-line), interquartile range (box), and 1.5*×* IQR (whiskers); each point is one sample. The best-performing setting is highlighted in bold. Asterisks denote significant differences from the best-performing setting (paired Wilcoxon signed-rank test; \**p <* 0.05, \*\**p <* 0.01, \*\*\**p <* 0.001, ns, not significant).

**Extended Data Fig. 7:**
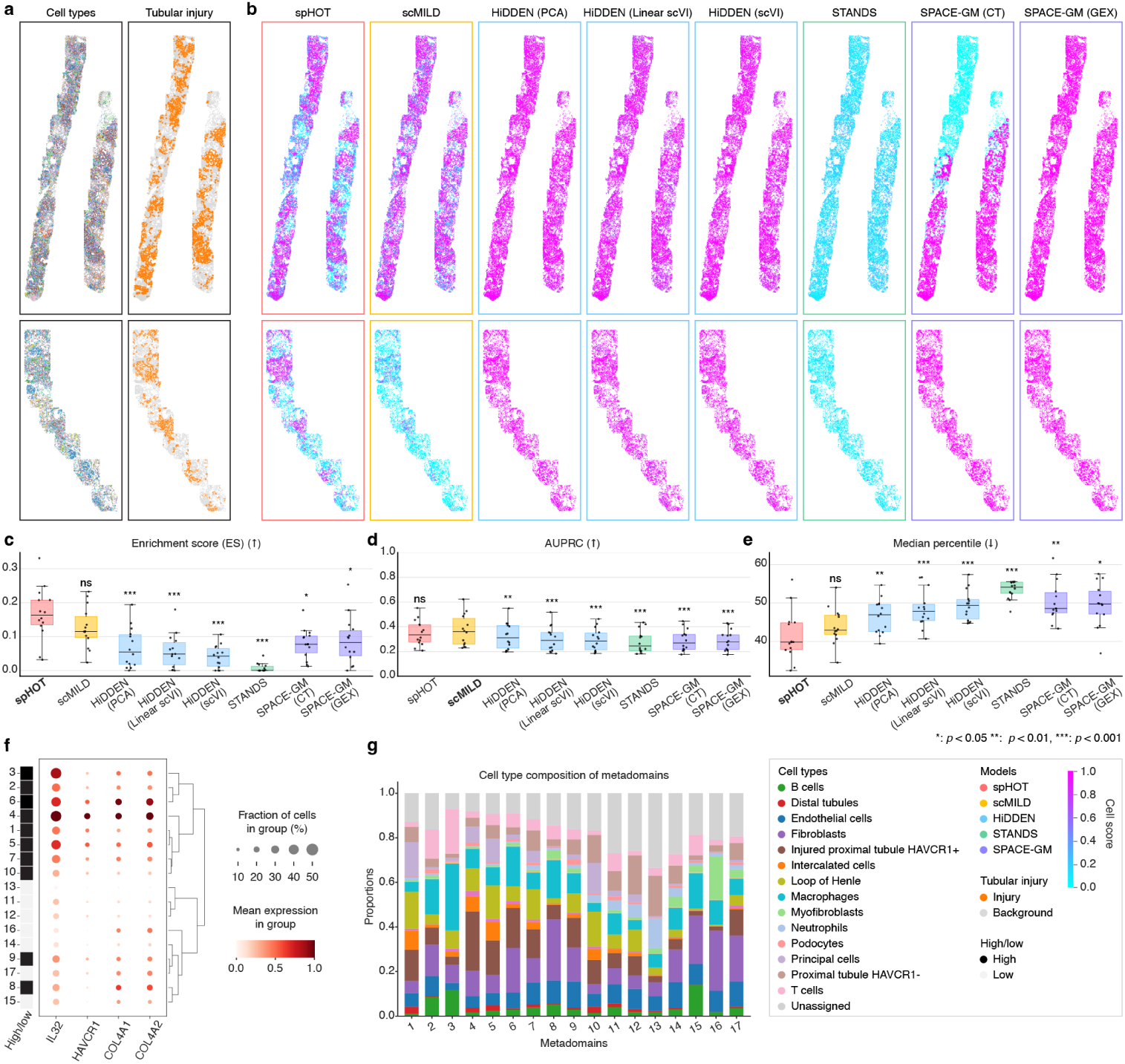
Benchmarking and downstream characterization for tubular injury region discovery in diabetic kidney disease. **a** Two representative samples showing cell type annotations (left) and ground truth tubular injury regions (right). **b** Continuous heatmaps of normalized cell scores within [0, 1] range for all methods. Box outlines indicate method type. Upper and lower rows correspond to the same representative samples as in **a**. **c**–**e** Benchmarking performance on tubular injury detection across 14 test-set case samples. Box plots show **c** ES; **d** AUPRC; **e** median ground truth percentile. **f** Dot plot of DKD injury marker expression across 17 metadomains, hierarchically clustered by expression profile. Dot size indicates the fraction of cells expressing each gene; color intensity indicates normalized mean expression level. Left axis annotations denote high/low-scoring metadomains as determined by majority voting across three cross-validation folds. **g** Cell type composition of all 17 metadomains, displayed as stacked bar plots. Colors correspond to annotated DKD cell types. Box plots show the median (centerline), interquartile range (box), and 1.5*×* IQR (whiskers); each point is one sample. The best-performing method is highlighted in bold. Asterisks denote significant differences from the best-performing method (paired Wilcoxon signed-rank test; \**p <* 0.05, \*\**p <* 0.01, \*\*\**p <* 0.001, ns, not significant).

**Extended Data Fig. 8:**
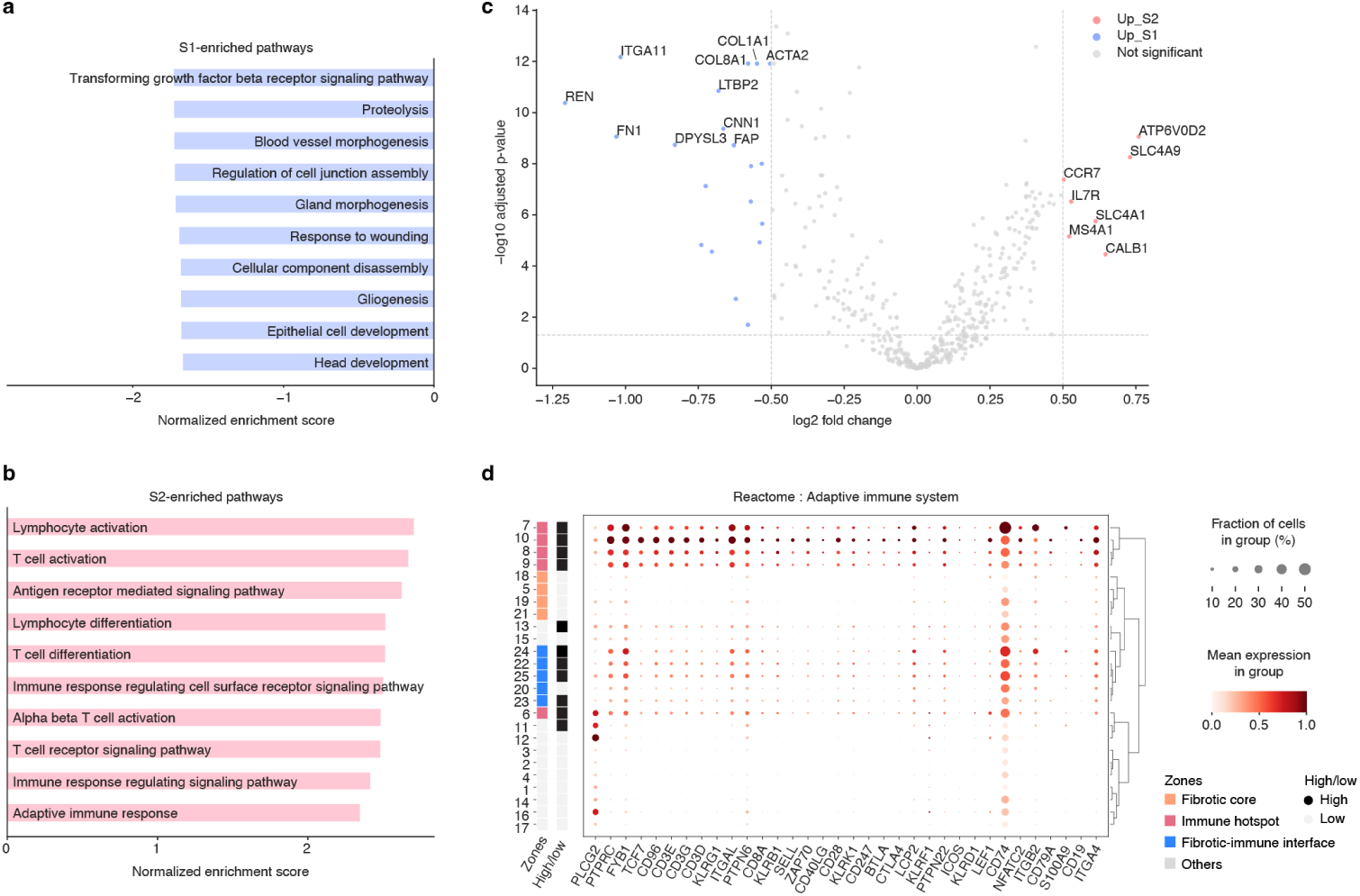
Pseudobulk differential expression analysis between S1-high and S2-high ANCA-GN cells. **a**–**b** Gene Ontology Biological Process (GO-BP) pathway enrichment results for S1-high-enriched pathways (**a**) and S2-high-enriched pathways (**b**), derived from preranked GSEA on pseudobulk differential expression between S1-high and S2-high cells. Bar plot shows normalized enrichment scores (NES) for top 10 significantly enriched pathways (FDR *<* 0.05). **c** Volcano plot of pseudobulk DE analysis. Top 10 significant gene symbols are denoted for both S1-upregulated and S2-upregulated genes. **d** Dot plot of Reactome adaptive immune system pathway genes across 25 S2 metadomains, hierarchically clustered by expression profile. Dot size indicates the fraction of cells expressing each gene; color intensity indicates normalized mean expression. Left axis annotations indicate zone assignment and right axis annotations show high/low metadomain classification determined by majority voting across cross-validation folds.

**Extended Data Fig. 9:**
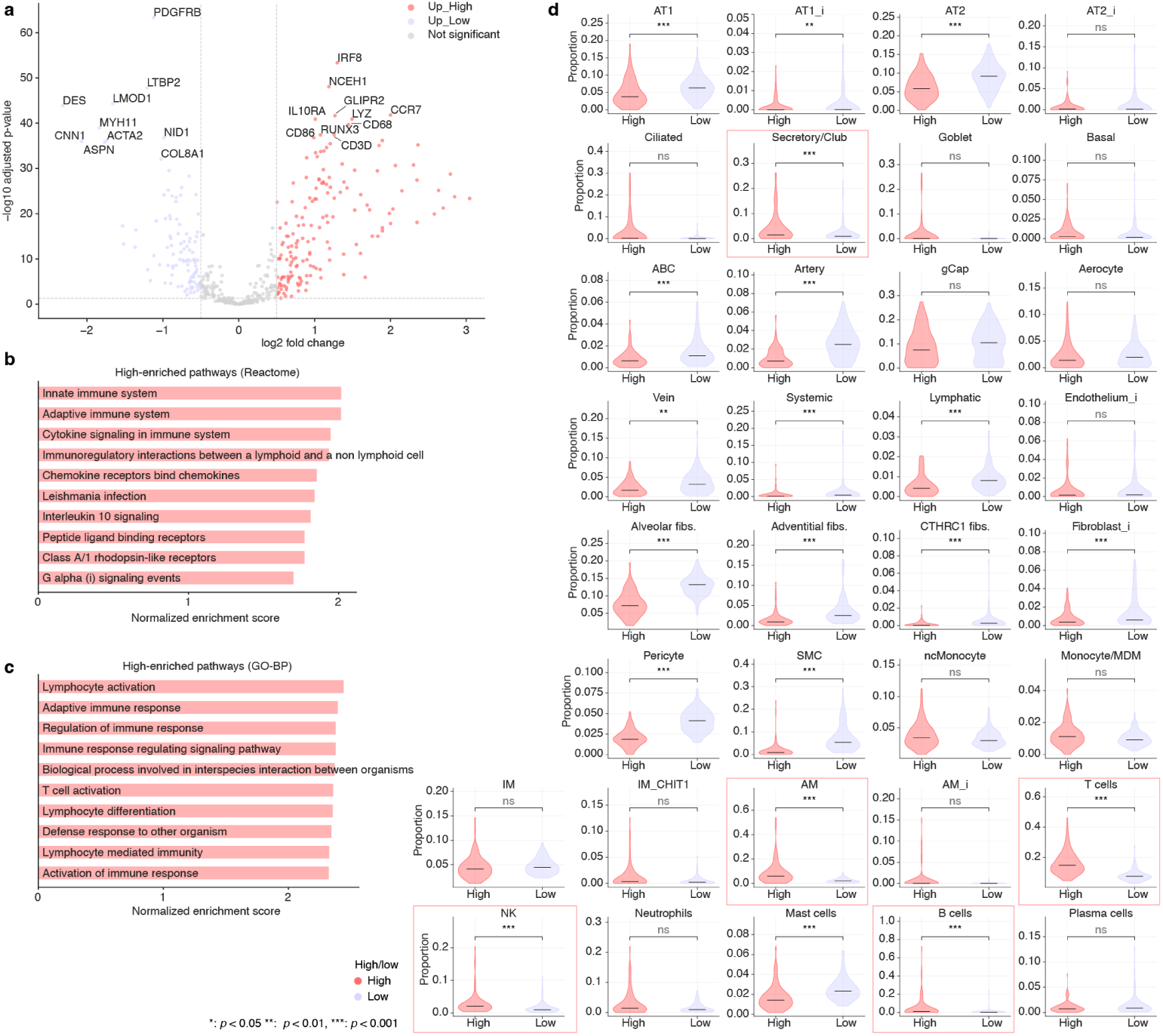
Transcriptomic and cellular characterization of high cells in COPD. **a** Volcano plot of pseudobulk DE analysis. Top 10 significant gene symbols are denoted for both high-upregulated and low-upregulated genes. **b**–**c** Reactome pathway enrichment results for high-enriched pathways (**b**) and Gene Ontology Biological Process (GO-BP) pathway enrichment results for high-enriched pathways (**c**), derived from preranked GSEA on pseudobulk differential expression between low and high cells. Barplot shows normalized enrichment scores (NES) for top significantly enriched pathways (FDR *<* 0.05). **d** Per-core cell type proportions in high-versus low-cells for all 34 annotated cell types (two-sided Mann-Whitney U test, Benjamini-Hochberg corrected). High cells are significantly enriched for immune populations including T cells, B cells, NK cells, and alveolar macrophages (boxed). Violins show the distribution across cores; horizontal lines denote medians. Individual data points are omitted. \**p <* 0.05, \*\**p <* 0.01, \*\*\**p <* 0.001, ns, not significant.

**Extended Data Table. 1:** Sample-level phenotype classification performance on simulated datasets.

| Model type | Model | Dataset |  |  |  |  |
| --- | --- | --- | --- | --- | --- | --- |
|  |  | Circular | Circular + Vessel-like | Vessel-like | CC | CCP |
| Single-cell-based | ProtoCell4P | <u>0.5140</u> | 0.4240 | 0.7120 | 0.9240 | **1.0000** |
|  | (prototype = 8) | (0.1191) | (0.3091) | (0.2186) | (0.1699) | (0.0000) |
|  | Hier-MIL | 0.4875 | 0.4000 | 0.6000 | 0.5000 | **1.0000** |
|  |  | (0.2978) | (0.1746) | (0.2746) | (0.3094) | (0.0000) |
|  | scMILD | 0.3625 | 0.4562 | 0.4188 | 0.7125 | 0.8000 |
|  |  | (0.2313) | (0.2313) | (0.1318) | (0.3175) | (0.4472) |
| Spatial-based | SPACE-GM (CT) | **1.0000** | 0.9125 | 0.9500 | 0.3750 | 0.4000 |
|  |  | (0.0000) | (0.1750) | (0.1000) | (0.4610) | (0.4899) |
|  | SPACE-GM (GEX) | **1.0000** | **0.9875** | 0.8875 | 0.8750 | 0.6875 |
|  |  | (0.0000) | (0.0250) | (0.2250) | (0.1118) | (0.3771) |
|  | spHOT | **1.0000** | <u>0.9250</u> | **1.0000** | **1.0000** | <u>0.9875</u> |
|  |  | (0.0000) | (0.0919) | (0.0000) | (0.0000) | (0.0250) |

Values are mean AUROC across five cross-validation folds, with standard deviation in parentheses. Bold indicates the best performance and underline the second best within each dataset. CT, cell type; GEX, gene expression; CC, circular and composition altered; CCP, circular and composition altered with perturbation.

## References

[1] Longo, S. K., Guo, M. G., Ji, A. L. & Khavari, P. A. Integrating single-cell and spatial transcriptomics to elucidate intercellular tissue dynamics. Nature Reviews Genetics 22, 627–644 (2021).

[2] Torres-Cano, A. et al. Spatially organized cellular communities shape functional tissue architecture in the pancreas. Science Advances 11, eadx5791 (2025).

[3] Xuanyuan, Q. et al. Multimodal spatial transcriptomic characterization of mouse kidney injury and repair. Nature Communications 16, 7567 (2025).

[4] Zhou, Y. & Glass, C. K. Microglia networks within the tapestry of alzheimer’s disease through spatial transcriptomics. Molecular Neurodegeneration 20, 102 (2025).

[5] Pelka, K. et al. Spatially organized multicellular immune hubs in human colorectal cancer. Cell 184, 4734–4752.e20 (2021).

[6] Cho, K. S. et al. Pan-cancer spatial atlas of tertiary lymphoid structures. Science 392, eadz2742 (2026).

[7] Marx, V. Method of the Year: spatially resolved transcriptomics. Nature Methods 18, 9–14 (2021).

[8] Moses, L. & Pachter, L. Museum of spatial transcriptomics. Nature Methods 19, 534–546 (2022).

[9] He, S. et al. High-plex imaging of RNA and proteins at subcellular resolution in fixed tissue by spatial molecular imaging. Nature Biotechnology 40, 1794–1806 (2022).

[10] Janesick, A. et al. High resolution mapping of the tumor microenvironment using integrated single-cell, spatial and in situ analysis. Nature Communications 14, 8353 (2023).

[11] Li, J., et al. Pan-cancer analysis of spatial transcriptomics reveals heterogeneous tumor spatial microenvironment. Cell Reports Medicine 7 (2026).

[12] Li, Y. et al. Spatial transcriptomics atlas of inflammatory bowel disease to guide implementation in research consortiums and clinical trials. Nature Communications 17, 5808 (2026).

[13] Hu, J. et al. SpaGCN: Integrating gene expression, spatial location and histology to identify spatial domains and spatially variable genes by graph convolutional network. Nature Methods 18, 1342–1351 (2021).

[14] Dong, K. & Zhang, S. Deciphering spatial domains from spatially resolved transcriptomics with an adaptive graph attention auto-encoder. Nature Communications 13, 1739 (2022).

[15] Ren, H., Walker, B. L., Cang, Z. & Nie, Q. Identifying multicellular spatiotemporal organization of cells with SpaceFlow. Nature Communications 13, 4076 (2022).

[16] Xu, C. et al. DeepST: identifying spatial domains in spatial transcriptomics by deep learning. Nucleic Acids Research 50, e131–e131 (2022).

[17] Long, Y. et al. Spatially informed clustering, integration, and deconvolution of spatial transcriptomics with GraphST. Nature Communications 14, 1155 (2023).

[18] Xu, H. et al. Unsupervised spatially embedded deep representation of spatial transcriptomics. Genome Medicine 16, 12 (2024).

[19] Zhu, B. et al. CellLENS enables cross-domain information fusion for enhanced cell population delineation in single-cell spatial omics data. Nature Immunology 26, 963–974 (2025).

[20] Duan, B., Chen, S., Cheng, X. & Liu, Q. Multi-slice spatial transcriptome domain analysis with SpaDo. Genome Biology 25, 73 (2024).

[21] Qian, J. et al. Identification and characterization of cell niches in tissue from spatial omics data at single-cell resolution. Nature Communications 16, 1693 (2025).

[22] Varrone, M., Tavernari, D., Santamaria-Martínez, A., Walsh, L. A. & Ciriello, G. CellCharter reveals spatial cell niches associated with tissue remodeling and cell plasticity. Nature Genetics 56, 74–84 (2024).

[23] Tejada-Lapuerta, A. et al. Nicheformer: a foundation model for single-cell and spatial omics. Nature Methods 22, 2525–2538 (2025).

[24] Blampey, Q. et al. Novae: a graph-based foundation model for spatial transcriptomics data. Nature Methods 22, 2539–2550 (2025).

[25] Goeva, A. et al. HiDDEN: a machine learning method for detection of disease-relevant populations in case-control single-cell transcriptomics data. Nature Communications 15, 9468 (2024).

[26] Do, C. & Lähdesmäki, H. Incorporating hierarchical information into multiple instance learning for patient phenotype prediction with single-cell RNA-sequencing data. Bioinformatics 41, i96–i104 (2025).

[27] Craig, E. et al. Annotation-free discovery of disease-relevant cells in single-cell datasets. Science Advances 11, eadv5019 (2025).

[28] Jeong, K., Choi, J. & Kim, K. scMILD: Single-cell multiple instance learning for sample classification and associated subpopulation discovery. iScience 29 (2026).

[29] Wu, Z. et al. Graph deep learning for the characterization of tumour microenvironments from spatial protein profiles in tissue specimens. Nature Biomedical Engineering 6, 1435–1448 (2022).

[30] Xu, K. et al. Detecting anomalous anatomic regions in spatial transcriptomics with STANDS. Nature Communications 15, 8223 (2024).

[31] Thorsson, J., Zhao, Y., Villablanca, E. J. & Engblom, C. Uncovering Immune Niches in Health and Disease Using Spatial Transcriptomics. European Journal of Immunology 56, e70185 (2026).

[32] Shaban, M. et al. MAPS: pathologist-level cell type annotation from tissue images through machine learning. Nature Communications 15, 28 (2024).

[33] Ilse, M., Tomczak, J. & Welling, M. *Attention-based Deep Multiple Instance Learning*, Vol. 80 of *Proceedings of Machine Learning Research*, 2127–2136 (PMLR, 2018).

[34] Moran, P. A. P. Notes on Continuous Stochastic Phenomena. Biometrika 37, 17–23 (1950).

[35] Geary, R. C. The Contiguity Ratio and Statistical Mapping. The Incorporated Statistician 5, 115–146 (1954).

[36] Qian, J. et al. Simulating multiple variability in spatially resolved transcriptomics with scCube. Nature Communications 15, 5021 (2024).

[37] Olivas, A. D. et al. Histopathologic Evaluation and Single-cell Spatial Transcriptomics of the Colon Reveal Cellular and Molecular Abnormalities Linked to J-Pouch Failure in Patients With Inflammatory Bowel Disease. Cellular and Molecular Gastroenterology and Hepatology 19 (2025).

[38] Xiong, G., Bekiranov, S. & Zhang, A. ProtoCell4P: an explainable prototype-based neural network for patient classification using single-cell RNA-seq. Bioinformatics 39, btad493 (2023).

[39] Vannan, A. et al. Spatial transcriptomics identifies molecular niche dysregulation associated with distal lung remodeling in pulmonary fibrosis. Nature Genetics 57, 647–658 (2025).

[40] Morse, C. et al. Proliferating SPP1/MERTK-expressing macrophages in idiopathic pulmonary fibrosis. European Respiratory Journal 54, 1802441 (2019).

[41] Mayr, C. H. et al. Spatial transcriptomic characterization of pathologic niches in IPF. Science Advances 10, eadl5473 (2024).

[42] Meadows, K. et al. Spatial transcriptomics identifies IL-32 as a lipid droplet-associated cytokine linked to tubular injury in human diabetic kidney disease. Inflammation Research 75, 33 (2026).

[43] Sultana, Z. et al. Spatiotemporal interaction of immune and renal cells controls glomerular crescent formation in autoimmune kidney disease. Nature Immunology 26, 1977–1988 (2025).

[44] Zhang, Y. et al. Aberrant cellular communities underlying disease heterogeneity in chronic obstructive pulmonary disease. Nature Genetics 58, 376–391 (2026).

[45] Virtanen, P. et al. SciPy 1.0: Fundamental Algorithms for Scientific Computing in Python. Nature Methods 17, 261–272 (2020).

[46] Ranasinghe, K., Naseer, M., Hayat, M., Khan, S. & Khan, F. S. Orthogonal Projection Loss, 12333–12343 (2021).

[47] Gayoso, A. et al. A Python library for probabilistic analysis of single-cell omics data. Nature Biotechnology 40, 163–166 (2022).

[48] Subramanian, A. et al. Gene set enrichment analysis: A knowledge-based approach for interpreting genome-wide expression profiles. Proceedings of the National Academy of Sciences 102, 15545–15550 (2005).

[49] Saito, T. & Rehmsmeier, M. The Precision-Recall Plot Is More Informative than the ROC Plot When Evaluating Binary Classifiers on Imbalanced Datasets. PLOS ONE 10, e0118432 (2015).

[50] Bradley, A. P. The Use of the Area under the ROC Curve in the Evaluation of Machine Learning Algorithms. Pattern Recognition 30, 1145–1159 (1997).

[51] Wolf, F. A., Angerer, P. & Theis, F. J. SCANPY: large-scale single-cell gene expression data analysis. Genome Biology 19, 15 (2018).

[52] Ellson, J., Gansner, E., Koutsofios, L., North, S. C. & Woodhull, G. Mutzel, P., Jünger, M. & Leipert, S. (eds) Graphviz— Open Source Graph Drawing Tools. (eds Mutzel, P., Jünger, M. & Leipert, S.) Graph Drawing, 483–484 (Springer Berlin Heidelberg, 2002).

[53] Muzellec, B., Teleńczuk, M., Cabeli, V. & Andreux, M. Pydeseq2: a python package for bulk rna-seq differential expression analysis. Bioinformatics 39, btad547 (2023).

[54] Fang, Z., Liu, X. & Peltz, G. GSEApy: a comprehensive package for performing gene set enrichment analysis in Python. Bioinformatics 39, btac757 (2023).

[55] Ragueneau, E. et al. The Reactome Knowledgebase 2026. Nucleic Acids Research 54, D673–D681 (2026).

[56] Ashburner, M. et al. Gene Ontology: tool for the unification of biology. Nature Genetics 25, 25–29 (2000).

[57] The Gene Ontology, C. The Gene Ontology knowledgebase in 2026. Nucleic Acids Research 54, D1779–D1792 (2026).

[58] Liberzon, A. et al. Molecular signatures database (MSigDB) 3.0. Bioinformatics 27, 1739–1740 (2011).

[59] Palla, G. et al. Squidpy: a scalable framework for spatial omics analysis. Nature Methods 19, 171–178 (2022).

[60] Kim, H., Kim, D., Jung, S., Lee, S. & Kim, K. spHOT Simulation Dataset: scCube-generated spatial transcriptomics benchmarking data for spatial biomarker discovery (2026). URL 10.5281/zenodo.21156231.

