## Supplementary Figures 1-2 for "Phenotype-associated spatial biomarker discovery in spatial transcriptomics with spHOT"

### 1 Supplementary Figure

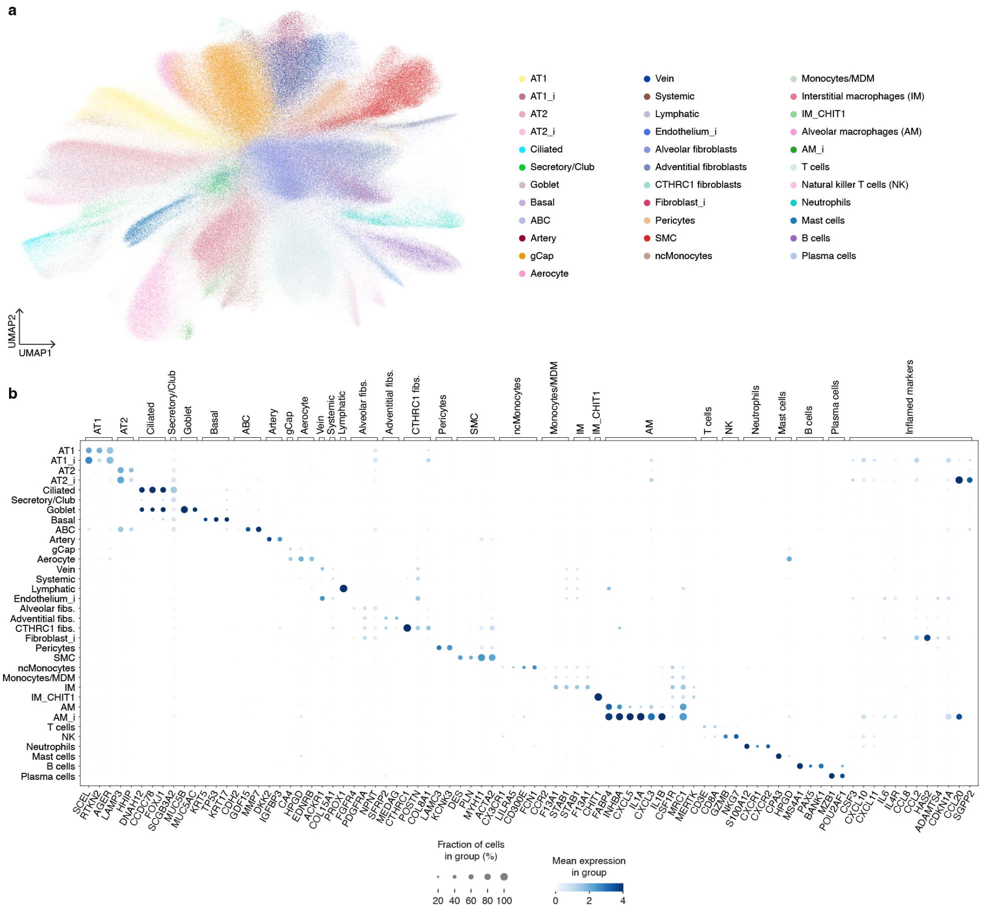


**Supplementary Fig. 1: Identification cell types in Xenium COPD dataset. a** UMAP visualization of total 816,245 quality controlled cells, colored by 34 annotated cell types. **b** Dot plot of canonical markers identifying cell types. Dot size denotes fraction of cells in group, color intensity indicates mean normalized expression.

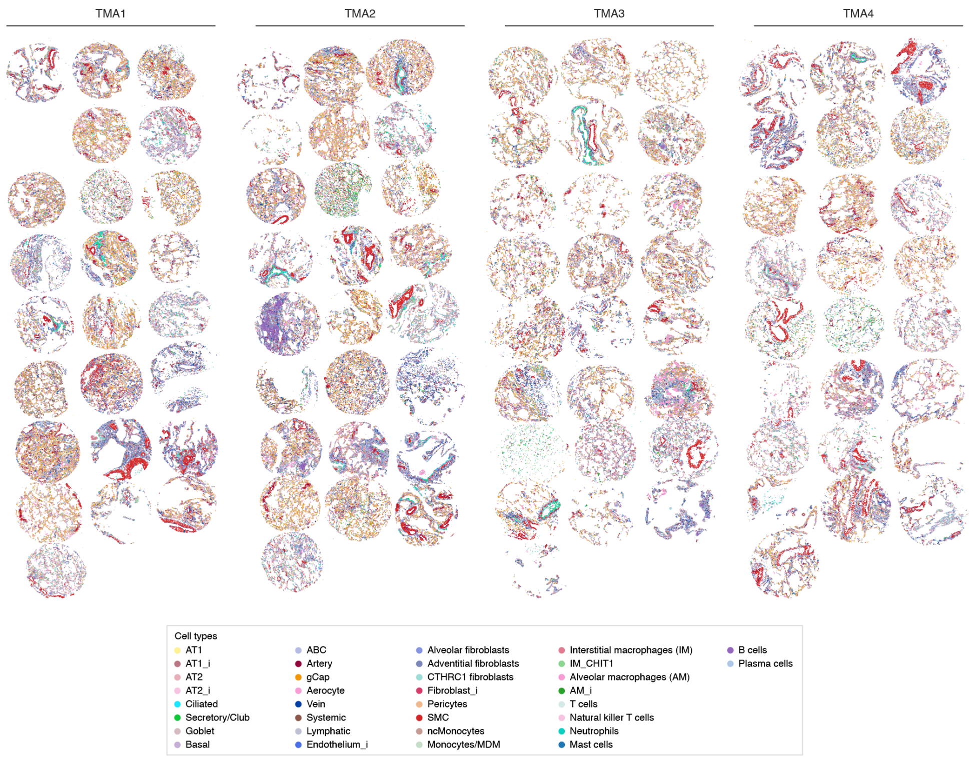


**Supplementary Fig. 2: Cell type maps of COPD tissue microarray slides.** All 99 TMA cores are color-coded based on their annotated cell types.
